# The genomic and epigenomic landscapes of hemizygous genes across crops with contrasting reproductive systems

**DOI:** 10.1101/2024.05.24.595664

**Authors:** Yanling Peng, Yiwen Wang, Yuting Liu, Xinyue Fang, Lin Cheng, Qiming Long, Dalu Su, Tianhao Zhang, Xiaoya Shi, Xiaodong Xu, Qi Xu, Nan Wang, Fan Zhang, Zhongjie Liu, Hua Xiao, Jin Yao, Ling Tian, Wei Hu, Songbi Chen, Haibo Wang, Sanwen Huang, Brandon S. Gaut, Yongfeng Zhou

## Abstract

Hemizygous genes, which present on only one of the two homologous chromosomes of diploid organisms, have been mainly studied in the context of sex chromosomes and sex-linked genes. However, structural variants (SVs), such as a deletion/insertion of one allele, can lead to hemizygous genes, a phenomenon largely unexplored in plants. Here, we investigated the genomic and epigenomic landscapes of hemizygous genes across 22 genomes with varying propagation histories: eleven clonal lineages, seven outcrossed samples and four inbred and putatively homozygous genomes. We identified SVs leading to genic hemizygosity. As expected, very few genes (0.01%-1.2%) were hemizygous in the homozygous genomes, representing negative controls. Hemizygosity was appreciable among outcrossed lineages, averaging 8.7% of genes, but consistently elevated for the clonal samples at 13.8% genes, likely reflecting heterozygous SVs accumulation during clonal propagation. Compared to diploid genes, hemizygous genes were more often situated in centromeric than telomeric regions and underlying weaker purifying selection. They also had reduced levels of expression, averaging ∼20% of the expression levels of diploid genes, violating the evolutionary model of dosage compensation. We also detected higher DNA methylation levels, in hemizygous genes and transposable elements, which may contribute to the reduced expression of hemizygous genes. Finally, expression profiles showed that hemizygous genes were more specifically expressed in contexts related to fruit development, organ differentiation and stress responses. Overall, hemizygous genes accumulate in clonally propagated lineages and display distinct genetic, and epigenetic features compared to diploid genes, shedding new insights on genetic studies and breeding programs of clonal crops.

## Introduction

Hemizygous genes are present on only one of the two homologous chromosomes of a diploid organism (1–3). The most prominent examples of hemizygous genes are on the sex chromosomes of male mammals (XY) or female birds (ZW) (1–3). Similar sets of hemizygous genes are present in the sex-linked regions of dioecious plants with X/Y sex determination, such as palms (4, 5), asparagus (6, 7), kiwifruit (8, 9). Numerous studies have focused on the evolution, gene expression, and epigenetic regulation of sex-linked hemizygous genes compared to diploid genes (2, 10–15). For example, genomic studies in mammalian males have consistently revealed lower mutation rates and more efficient selection in sex-linked hemizygous genes than diploid genes (10), due in part to the fact that hemizygosity uncovers recessive alleles and makes them visible to selection (16). Similar studies have generally shown that the ratio of sex-linked to diploid (X:AA) (where X represents sex-linked genes and AA represents autosome genes) gene expression is ∼0.5 in animals and plants (11, 17). This ratio is inconsistent with the hypothesis that dosage compensation re-equalizes male and female expression to restore XY male expression back to its ancestral level (17).

Interestingly, estimated expression levels of XY sex chromosome alleles in males show an overall trend of reduced expression of Y-linked alleles relative to X-linked alleles in both animals and plants (12–15). Some of these expression effects are mediated by epigenetic marks, including histone modifications and DNA methylation (18–21). For example, the male-specific region of the papaya Y chromosome is associated with knob-like heterochromatic structures that are heavily methylated. This suggests that DNA methylation has played a role in the evolution of this Y chromosome (18). These observations indicate that sex-linked hemizygous genes often have distinct epigenetic and regulatory features. In addition to sex-linked regions, the absence of one paired allele is frequently observed in non-sex-linked regions of homologous chromosomes in diploid plants, leading to a significant presence of hemizygous genes (22). However, the extent and function of these genes remain largely uncharacterized.

Here, we study hemizygous genes across a collection of plant genomes with contrasting propagation histories. The identification of these hemizygous (or haploid) genes in diploid plant genomes has become feasible with the emergence of long-read sequencing technologies and the advancements in assembly algorithms. Precise genome assemblies facilitate the identification of structural variants (SVs) in heterozygous diploid plant genomes, thereby permit genome-wide identification of hemizygous genes caused by SVs (22–26). For example, by remapping long-reads to a reference genome assembly, it has been inferred that ∼13.5% and ∼15% of genes are hemizygous in two clonal grapevine (*Vitis vinifera* ssp. *vinifera*) cultivars (22). This high value may in part reflect unique features of long-term clonal lineages, because recessive deleterious mutations are expected to accumulate in these lineages (22, 27). Nonetheless, hemizygosity is not limited solely to clonal lineages, because ∼8.89% and ∼4% of genes are estimated to be hemizygous in an outcrossing wild rice species (*Oryza longistaminata*) and in avocado (*Persea americana*), respectively (28, 29). In contrast, as expected, only a few genes have been inferred to be hemizygous in inbred, self-fertilized accessions. For example, only 0.73% and 0.35% of genes were inferred as hemizygous in rice cultivars Nipponbare and 93-11, respectively (28).

The extent of hemizygosity in plant genomes has only begun to be appreciated, largely because genome projects have historically focused on self-fertilized or homozygous materials (22). As a result, there is currently little information about natural variation in the number of hemizygous genes, about potential correlations between hemizygosity and life history traits, such as reproductive systems and historical population sizes. There is also limited information about the evolutionary dynamics and putative functions of hemizygous genes. However, we are aware that hemizygosity can affect function. For example, white berry color in grapevines is related to a complex series of mutations, which includes hemizygosity of a large genomic region (30). In this case, the key feature of hemizygosity is that it uncovered a recessive, non-functional allele that interrupts anthocyanin biosynthesis. The *ClpC*-like gene in *Citrus clementina* is also hemizygous and exhibits lower expression compared to the wild-type plants, resulting in a reduced chlorophyll a/b ratio in green tissues (31). The *MdACT7* gene, located within a 2.8-Mb heterozygous deletion, is expressed at lower levels in *Malus domestica* "AGala" compared to "Gala", causing delayed fruit maturation in the former (32). Functional effects are not limited to plants; hemizygous deletions in human autosomes are often associated with diseases and cancer. These deletions are often accompanied by decreased gene expression and increased DNA methylation, as illustrated by the examples of genes like *RUNX3*, *KLF4*, and *TP53* (33–35).

Thus far, the extent of hemizygosity has only been estimated in a handful of plant genomes, and there have been no accompanying genome-wide analyses of hemizygous gene function and epigenetic states. In this study, we build or gather haplotype-resolved genomes and primary assemblies for 22 genome samples, all based on PacBio HiFi data, and subsequently identify SVs that define hemizygous genes. The 22 samples represent a range of reproductive histories, including eleven clonally propagated plants representing grapevines (*Vitis vinifera*), apples (*Malus domestica*) and cassava (*Manihot esculenta*). Since extensive hemizygosity may be elevated in clonal lineages, we have also included comparisons from seven outcrossing samples -including wild grapevines, wild apples, and one wild rice species (*Oryza rufipogon*) - and four inbred cultivars (*SI Appendix,* Table S1). To compare results across samples, we have assembled PacBio HiFi sequencing data for each genome and identified SVs that define hemizygous genes.

Given the identification of hemizygous genes, along with the availability of transcriptomic and epigenomic data for a subset of the samples, we ask the following questions: (i) Are hemizygous genes widespread in diploid plant genomes, or are they particularly abundant in clonal lineages? (ii) Do hemizygous genes have distinct sequences and evolutionary features compared to diploid genes? For example, are they enriched for specific biological processes? If they are, are they expressed at half the average expression levels of diploid genes? (iii) Hemizygous regions can include genes as well as other sequence features, like transposable elements (TEs). How extensive are hemizygous TEs, and do they have detectable correlations with the expression of nearby diploid genes? Finally, (iv) Do hemizygous genes exhibit distinct epigenetic patterns when compared to diploid genes? If they do, is expression related to these epigenetic effects? By addressing these questions, our goal is to further understand the evolutionary and functional consequences of genic hemizygosity. Ultimately this knowledge will be beneficial for understanding the genetics, breeding, and evolution of plants with heterozygous genomes.

## Results

### The prevalence of hemizygous genes in clonal plant genomes

To identify hemizygous genes, we either built or gathered haplotype-resolved genome assemblies for 11 clonal and seven outcrossing plants, as well as primary genome assemblies for four inbred samples (*SI Appendix*, Table S1). The four inbred samples, representing grapevine, tomato, and rice, were included as a control, because hemizygosity should be near zero in these lineages. All 22 surveyed genomes were assembled with PacBio HiFi data (*SI Appendix,* Table S2). Among the assemblies, three *Vitis* genomes were newly generated based on PacBio HiFi and Hi-C data, and haplotypes were resolved with ultra-long Oxford Nanopore Technologies (ONT) data (> 100kb) for gap-closing (for Chardonnay) (*SI Appendix,* Fig. S1). The three new haplotype-resolved genome assemblies were anchored to 38 chromosomes and highly contiguous, with scaffold N50 sizes ranging from 25.1 to 26.3 Mb and Benchmarking Universal Single-Copy Orthologs (BUSCO) completeness scores of 97.3% to 98.5% (*SI Appendix,* Table S3). The data and reference assemblies for the remaining 19 accessions were retrieved from public repositories (*SI Appendix,* Table S2). The 19 assemblies were anchored to chromosome level, had scaffold N50 values of 25.1 to 67.6 Mb and BUSCO completeness scores of 93.0% to 99.2% (*SI Appendix,* Table S3).

Given these high-quality references, we identified hemizygous regions by remapping PacBio HiFi reads longer than 10 kb to genome assemblies, focusing on SVs longer than 50 bp. The SVs were identified using the Sniffles pipeline, followed by several filtering steps, including thresholds for quality and coverage (see Methods). Because the depth of coverage varied from ∼40×to 80× or higher across the 22 samples, we downsampled the PacBio HiFi data (reads longer than 10 kb) to 40×coverage before mapping to facilitate fair comparisons (*SI Appendix,* Fig. S4). Focusing on grapevine, this approach yielded between 59,015 and 61,066 heterozygous SVs (hSVs) in the three clonal samples. Of these, more than one-third (i.e., 26,845 to 27,982, 45.1%-45.8% of total hSVs) were heterozygous deletions (hDELs) relative to the reference (*SI Appendix,* Fig. S4A); among the remaining SV types, 13.8%-16.8% were classified as heterozygous breakends (hBND), duplications (hDUP) or inversions (hINV), while heterozygous insertions (hINS) accounted for 38.1%-40.4% of SVs. Similar patterns were observed in the remaining clonal samples (*SI Appendix,* Fig. S4A). We excluded the hINS class from further analyses because the reference lacks information about content within these SVs.

Even after excluding hINSs, our focus on hDELs, hBNDs, hDUPs, and hINVs revealed extensive hemizygosity within clonal plants compared to outcrossing and inbred lineages. In the three clonal genomes of grapevine, for example, hemizygous regions ranged from 76.0 to 124.6 Mb, corresponding to 15.4% to 25.6% of the total genome size (*SI Appendix,* Table S1). The genome-wide extent of hSVs in clonally propagated apples, potatoes and cassava was even higher than that of the grapevine varietals, perhaps reflecting their larger genome sizes (*SI Appendix,* Table S1) and potentially reflecting features of their reproductive and life histories, such as the duration of the clonal lineage.

The hemizygous genome proportion was substantially higher, averaging at ∼25%, for clonal samples compared to the outcrossing (∼16% on average) and inbred (< 1.1%) samples (*SI Appendix,* Table S1). The low proportion for the inbred samples suggest that our methods have low false positive errors – i.e., representing < 1.1 % of the genome. The differences in proportions of hemizygous genes were highly significant among clonal and outcrossing propagation histories (e.g., *P* < 0.05 between clonal and outcrossing samples; Mann-Whitney U-test). Although one must exercise caution with this comparison, as our samples were not phylogenetically independent, the data consistently indicated that clonally propagated lineages have elevated hemizygosity.

We further characterized hemizygous regions focusing on whether they contained complete genes or TEs (*SI Appendix,* Fig. S3). Focusing again on grapevines as an example, we detected between 5,131 and 6,573 hemizygous genes (representing between 12.6% and 16.8% of total genes) in the three clonal samples, with Thompson Seedless exhibiting the highest proportion (Fig. 1 and *SI Appendix,* Table S1). These genomes also contained numerous hemizygous TEs, between 174,850 and 194,536, representing 16.2% to 18.1% of annotated TEs (*SI Appendix,* Fig. S5 and Table S6), with Thompson Seedless again exhibiting the highest proportion. In contrast to the clonal samples, the outcrossing *Vitis* genomes had fewer hemizygous genes and TEs, with between 3,745 and 4,356 hemizygous genes representing 8.5% to 10.5% of total genes (*SI Appendix,* Table S1) and between 60,949 and 137,246 hemizygous TEs (between 5.4% and 12.2% of total TEs) (*SI Appendix*, Fig. S5 and Table S6). As expected, the inbred, nearly homozygous PN40024 *V. vinifera* genome had far fewer hemizygous genes and TEs, with 166 hemizygous genes (0.4% of annotated genes) and 4,828 hemizygous TEs (0.4% of total TEs). The analysis of nine *Malus*, one *Manihot*, two *Solanum*, and three *Oryza* accessions showed similar patterns (Fig. 1 and *SI Appendix*, Fig. S5, Table S1 and Table S6) – i.e., elevated gene and TE hemizygosity in clonal accessions. Altogether, these observations generalize previous findings, based on only a few genomes, which estimated that: i) ∼15% and 13.5% of genes are hemizygous in clonal grapevine cultivars (23, 27); ii) a lower but notable percentage of hemizygous genes in outcrossing plants (i.e., 8.89% hemizygous genes in *O. longistaminata* (28), 4% in avocado (29), and now between 5.9% and 11.7% in wild outcrossing samples (*SI Appendix*, Table S1); and iii) consistently low rates of genic hemizygosity (<1%) in putatively homozygous materials (28).

**Fig. 1.**
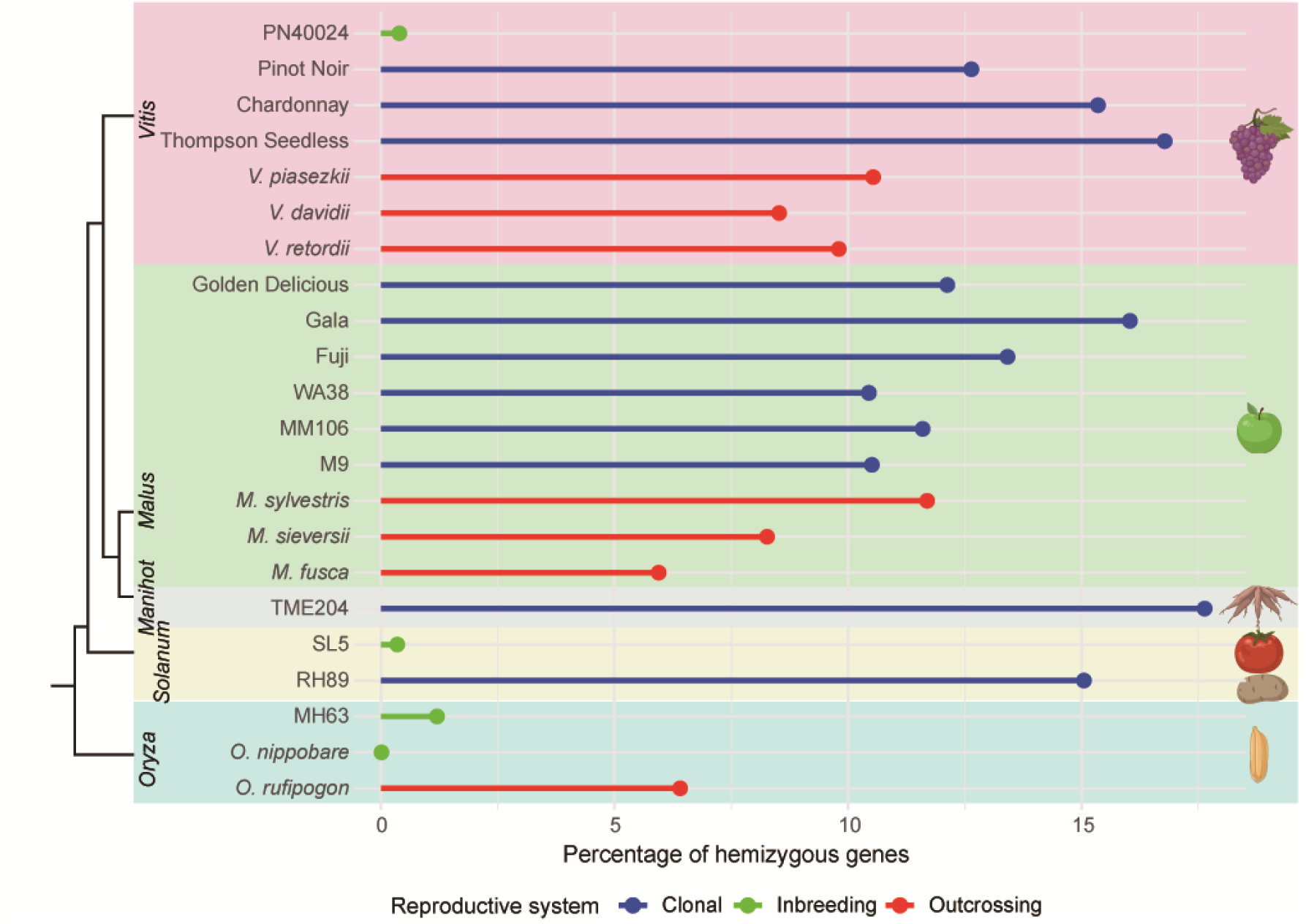
Proportion of hemizygous genes in crop genomes with contrasting reproductive systems. The phylogeny on the left is based on one previous study (95).

### Refinement of the set of hemizygous genes in *Vitis* assemblies

We anticipated that our observations based on Sniffles represented *bona fide* SVs. However, for downstream characterization, we deemed it critical to define a subset of genes with additional evidence supporting hemizygosity. For these analyses, we focused on the *Vitis* accessions because the genome and annotation pipeline were consistent (thereby limiting technical variation among groups or sequencing sites). Additionally, expression and epigenetic data were available for most samples, and the selected samples represent the range of reproductive histories considered in this study (*SI Appendix*, Table S1).

We identified a refined set of heterozygous SVs (hSVs) for the three clonal and three outcrossing *Vitis* samples (*SI Appendix*, Table S1) by aligning the haplotypes with each other, inferring hemizygous regions, and then taking the intersection with the Sniffles-based inferences (see Methods). This additional filter reduced the hemizygous gene set by ∼50% across the six samples to an average of 2,365 genes (range: 1,409 to 3,258) (*SI Appendix,* Fig. S6A). We focused on these genic sets for downstream analyses and contrasted them with diploid genes – i.e., genes without any evidence of an overlapping hSV. We did not include the inbred PN40024 sample in subsequent analyses, however, because it contained so few hemizygous genes.

Given this filtered set of hemizygous genes, we examined statistics such as the proportion of genes with a single exon, the number of exons per gene, exon length, and overall gene length (*SI Appendix,* Fig. S6B-E). The percentage of one-exon genes was significantly higher for hemizygous than diploid genes (Fisher exact test, *P* < 0.05; *SI Appendix,* Fig. S6B), but the average number of exons, exon length, and gene length of hemizygous genes were all significantly lower (*P* < 0.05, *SI Appendix,* Fig. S6C-S6E). These results were not unexpected, because one might naively expect that shorter genes have a higher probability of being encompassed by an SV event. It is also possible that the SV and alignment algorithms were biased toward identifying shorter genes, but we attempted to obviate potential biases by using only >10kb reads and adding an additional filter based on whole genome alignments. Altogether, these results suggest that hemizygous genes are shorter and structurally simpler than diploid genes.

### Evolutionary and functional properties of hemizygous genes

We then explored evolutionary and functional features of putatively hemizygous genes relative to diploid genes for the six *Vitis* samples, three clonal and three outcrossing wild (*SI Appendix*, Table S1). For instance, we analyzed the proportion of hemizygous genes in centromeric and telomeric regions, representing regions of lower and higher recombination, respectively. The proportion of hemizygous genes was higher in centromeric regions compared to diploid genes, whereas the opposite pattern was observed in telomeres (Fig. 2A). This suggests that hemizygous genes tend to be biased toward low recombination regions, where selection is less effective. We also measured nonsynonymous (*Ka*) and synonymous (*Ks*) substitution rates for each gene by comparing sequences to a paired outgroup (e.g., *Muscadinia rotundifolia*). In each of the samples, a lower percentage of hemizygous genes (2.5% across six genomes on average) were alignable to the outgroup than diploid genes (23.3% on average), suggesting that hemizygous genes either evolve more rapidly than diploid genes or are more dispensable (i.e., more often lost over evolutionary time). After aligning the available genes, the median *Ks* value in hemizygous genes was significantly lower than diploid genes in all six *Vitis* samples (Wilcoxon rank-sum test, *P <* 0.05; Fig. 2B), which could reflect that the alignable hemizygous genes are a conserved, biased subset. Nonetheless, these hemizygous genes have correspondingly higher median *Ka/Ks* values relative to diploid genes (Wilcoxon rank-sum test, *P <* 0.05; Fig. 2B), consistent with weaker purifying selection, lower mutation rates (as implied by lower *Ks* values) or some combination of these two processes.

**Fig. 2.**
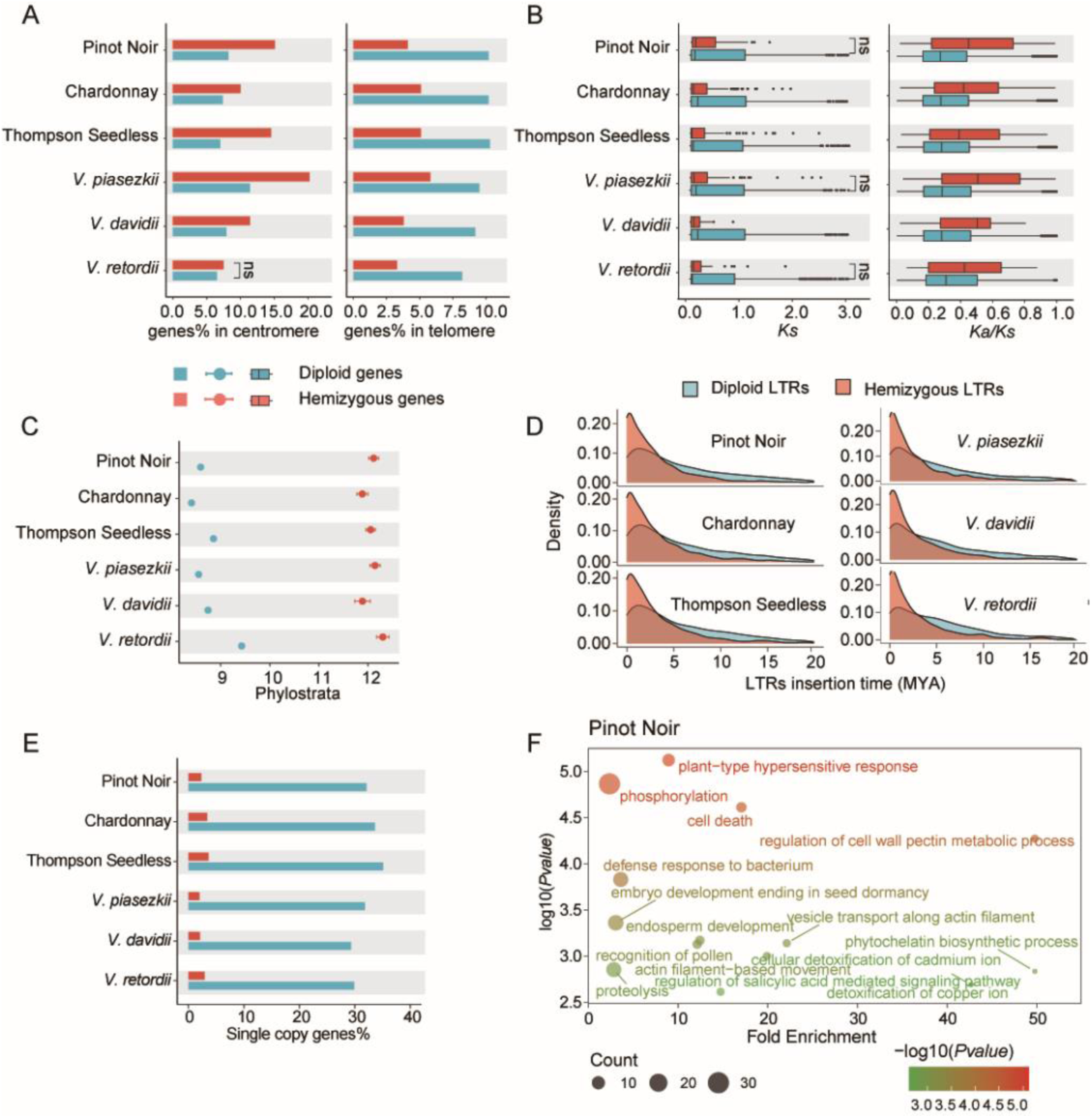
Characterization of hemizygous genes in clonal and outcrossing *Vitis* accessions. (A) Proportions of hemizygous and diploid genes located in centromeres (left) and telomeres (right), respectively. (B) *Ks* and *Ka/Ks* values comparing hemizygous and diploid genes; the line in the middle of the box represents the median, the edges of the box indicate first and 3rd quartiles, and the whiskers represent the range. (C) Average phylostrata levels based on phylostratigraphic analysis for both hemizygous and diploid genes, with larger phylostrata indicating younger genes. Error bars show 95% bootstrap-based confidence intervals. (D) Insertion times for hemizygous and diploid long terminal repeats (LTRs). (E) Proportion of single copy genes among total hemizygous and diploid genes. (F) The top 15 enriched biological processes for hemizygous genes in Pinot Noir. A Fisher’s exact test was conducted for (A) and (E), and a Wilcoxon rank-sum test was applied to (B), (C), and (D). Non-significant differences are indicated by "ns" (*P* > 0.05), while significant differences (*P* < 0.05) are shown without a mark.

We then examined the age of hemizygous genes by employing phylostratigraphic analyses (36). These analyses indicated that hemizygous genes originated more recently than diploid genes (Wilcoxon rank-sum test, *P <* 0.05; Fig. 2C). We also calculated the insertion times of hemizygous and diploid long terminal repeat (LTR) TEs, finding that hemizygous TEs are also evolutionary younger (Wilcoxon rank-sum test, *P <* 0.05; Fig. 2D).

We also investigated the proportion of single-copy and multi-copy genes in both hemizygous genes and diploid genes. We hypothesized that hemizygous genes were more likely to belong to a multi-gene family, because gene family membership can provide functional redundancies that make hemizygosity potentially less detrimental. To measure membership in gene families, we implemented a OrthoFinder-based pipeline (37). Our results supported our hypothesis, because we detected a lower proportion of single copy genes in hemizygous genes compared to that of diploid genes in all six plants (2.3% vs. 32.1% in Pinot Noir; 3.4% vs. 33.6% in Chardonnay, 3.6% vs. 35.1% in Thompson Seedless; 2.1% vs. 29.3% in *V. davidii*; 2.0% vs. 31.8% in *V. piasezkii*; 2.9% vs. 29.2% *V. retordii*; *P* < 0.05 for all cases; Fisher’s Exact Test) (Fig. 2E).

Finally, we investigated the possible biological processes of hemizygous genes in the six *Vitis* plants (Fig. 2F and *SI Appendix,* Fig. S8). In Pinot Noir, for example, the top 15 enriched Gene Ontology (GO) terms were involved in biological processes such as recognition of pollen, endosperm development, and defense response to bacterium. In Chardonnay, the top 15 enriched GO terms were related to recognition of pollen, floral organ development, defense response. In Thompson Seedless, the top 15 enriched GO terms were related to responses to abscisic acid, plant ovule development and immune response. In *V. piasezkii*, the top 15 enriched GO terms were involved in response to water deprivation, response to salt stress, defense response to bacterium. In *V. retordii*, the top 15 enriched GO terms were involved in response to recognition of pollen, endosperm development, and defense response to bacterium. In *V. davidii*, the top 15 enriched GO terms were involved in response to salt stress, recognition of pollen, and responses to abscisic acid. These results indicate that hemizygous genes can be involved in fundamental processes like reproduction and mitosis, but they are also consistently enriched for responses to biotic and abiotic stress.

### Unique expression patterns of hemizygous genes

A simple null expectation for hemizygous genes is that they are expressed at 50% of the average level of diploid genes. To explore this hypothesis, we amassed RNA-seq datasets generated across accessions, developmental stages (e.g., fruit development) and experimental regimes, such as stress treatments (*SI Appendix,* Table S8). The samples included 21 from different tissues/treatments in Pinot Noir, 30 belonging to 10 different tissues/treatments in Chardonnay, 20 from eight different tissues/treatments in Thompson Seedless, and 27 from 11 different tissues/treatments in *V. davidii*. We also generated three leaf samples from one control treatment in both *V. piasezkii* and *V. retordii* (*SI Appendix,* Table S8) and evaluated 29 samples from 12 different tissues/treatments for Golden Delicious apples (*SI Appendix,* Table S8). In total, we recovered 691 Gb paired-end and 74 Gb single-end Illumina reads from 168 samples representing six of our 22 samples. Our goals with these data were: i) to investigate the level of expression in hemizygous genes relative to diploid genes and ii) to determine whether hemizygous genes had patterns of expression consistent with contributions to development and other processes.

We first assessed whether hemizygous genes were expressed. Across taxa and individual RNA-seq samples, a significantly higher proportion of hemizygous genes had no evidence of expression relative to diploid genes (Fig. 3A, *SI Appendix,* Fig. S9A). For example, across all samples in Pinot Noir, 49.2% (1,334 out of 2,711 genes) of hemizygous genes had evidence for expression, but that proportion was 74.2% (26,309 out of 35,459 genes) for diploid genes (Chi-sq = 704.1, df = 1; *P* < 0.05). This trend held true in each tissue/treatment for all taxa – e.g., among the 21 RNA-seq tissues/treatments in Pinot Noir, hemizygous genes were expressed in lower proportions for all tissues/treatments. These results strongly suggest that hemizygous genes are enriched for pseudogenes or for a subset of genes that are expressed under fewer experimental and developmental conditions. However, not all hemizygous genes were pseudogenes; across all data samples, we detected expression for 49.2%, 66.4%, 44.5%, 24.7%, 52.7%, 29.4%, and 61.1% of hemizygous genes in Pinot Noir, Chardonnay, Thompson Seedless, *V. piasezkii*, *V. davidii,* and *V. retordii,* and Golden Delicious, respectively (Fig. 3A and *SI Appendix,* Fig. S9A).

**Fig. 3.**
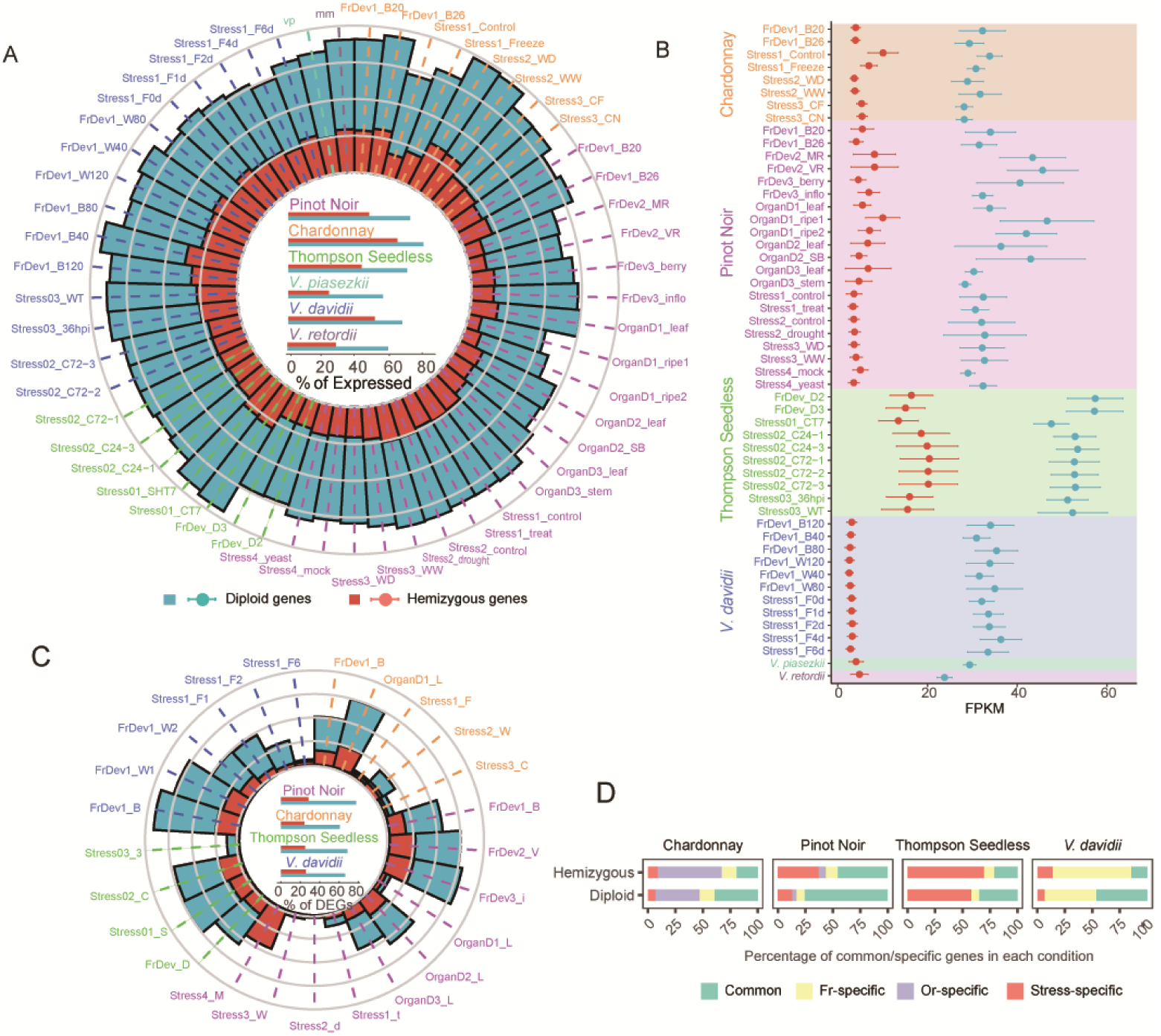
Transcriptomic landscapes of hemizygous and diploid genes in clonal and outcrossing *Vitis* samples. (A) The proportion of expressed hemizygous genes and diploid genes in each tissue/treatment and across all data (shown in the center) from the six *Vitis* samples. Each circle represents increments of 25%, with the innermost circle indicating 25% and the outmost circle is 100%. (B) Expression levels, measured as FPKM, for hemizygous and diploid genes in each tissue/treatment from the six *Vitis* samples. Error bars show 95% bootstrap-based confidence intervals. (C) The proportion of differentially expressed genes (%DEGs) for hemizygous and diploid genes for four *Vitis* samples (named in the center) that allow control-treatment contrasts. Each circle represents increments of 25%, with the innermost circle indicating 25% and the outermost circle indicating 100%. (D) The proportion of common and unique differentially expressed hemizygous and diploid genes among three processes, including fruit development (Fr), organ differentiation (Or), abiotic and biotic stress stimulus processes (Stress). A Fisher test was conducted for (A) and (C), while a Wilcoxon rank-sum test was applied to (B). Significant differences (*P* < 0.05) between diploid and hemizygous genes were observed in all cases.

We next investigated average levels of expression for the subset of genes with evidence for expression. Hemizygous genes were consistently expressed at significantly lower levels than diploid genes based on average expression across all tissues/treatments and within each tissue/treatment (Fig. 3B, *SI Appendix*, Fig. S9B). The hemizygous: diploid ratio of median expression was ∼0.07 for Pinot Noir and *V. davidii* but higher for other *Vitis* samples (range 0.14 to 0.21) and highest for Golden Delicious apples (0.32) (*SI Appendix,* Fig. S9C). Nonetheless, all these values were significantly (*P* < 0.05, Wilcoxon signed rank test) lower than the 50% expected if hemizygous alleles were expressed at similar levels to diploid alleles. These results imply that there is a diminution of gene expression associated with hemizygosity, and this diminution typically results in average expression levels of hemizygous genes being less than 50% of those of diploid genes (see Discussion).

We then examined patterns of hemizygous gene expression across specific functional categories that were explored in the original RNA-seq experiments. These categories included stages of fruit development, organ differentiation, and responses to biotic and abiotic stresses. First, we estimated the proportion of hemizygous genes that were differentially expressed across comparisons. Hemizygous genes generally had a lower proportion of differentially expressed genes than diploid genes (Fig. 3C and *SI Appendix,* Fig. S9D). For example, in Pinot Noir, 28.3% (378 out of 1,334) of hemizygous genes and 77.5% (20,394 out of 26,309) of diploid genes were differentially expressed across all paired comparisons (Chi-sq = 1712.0, df = 1; *P* < 0.05). The corresponding values for Chardonnay were 24.2% (454 out of 1,873) versus 60.7% (16,987 out of 27,978) (Chi-sq = 942.2, df = 1; *P* < 0.05), 24.8% (359 out of 1,450) versus 68.6% (16,168 out of 23,578) (Chi-sq = 1105.6, df = 1; *P* < 0.05) in Thompson Seedless, 25.5% (189 out of 742) versus 66.3% (18,468 out of 27,838) (Chi-sq = 515.9, df = 1; *P* < 0.05) in *V. davidii*, and 26.3% (366 out of 1,389) versus 53.4% (17,261 out of 32,313) (Chi-sq = 386.3, df = 1; *P* < 0.05) in Golden Delicious. Overall, fewer expressed hemizygous genes differed in expression during fruit development, organ differentiation or during abiotic and biotic stress.

We also explored expression specificity by detecting genes that were expressed in specific tissues or treatments. That is, we counted the number of genes that had significant evidence for expression in only one tissue/treatment of a paired comparison. Altogether, hemizygous genes had a higher proportion of tissue/treatment-specific genes than the diploid genes (*SI Appendix,* Fig. S10). Across 10 tissue/treatment comparisons in Pinot Noir, 31.0% to 50.0% of hemizygous genes were expressed in only one of the paired tissues or treatments, but these values were substantially lower for diploid genes (*SI Appendix,* Fig. S10). Similar patterns – i.e., more tissue/treatment-specific expression for hemizygous genes – were also found in Chardonnay, Thompson Seedless, *V. davidii* and Golden Delicious (*SI Appendix,* Fig. S10). Given these results, we anticipated that a lower proportion of hemizygous genes would be expressed across experiments investigating fruit development, organ differentiation, abiotic and stress; indeed, this was true for all relevant samples (Fig. 3D and *SI Appendix* Fig. S9E).

### The cis-regulatory effects of hemizygous and diploid TEs on gene expression

We then explored the cis-regulatory effects of TEs on gene expression. To do so, we used both the RepeatModeler and EDTA pipelines to identify TEs for the three clonal and three outcrossing *Vitis* samples, detecting from 337,284 to 502,618 TEs across the six accessions based on both pipelines (*SI Appendix,* Table S6). We then classified genes into four categories based on their proximity to annotated TEs. The four categories were: i) hemizygous genes with nearby TEs (i.e., within 2kb of either the 5’ or 3’ ends of genes), ii) hemizygous genes without nearby TEs, iii) diploid genes with nearby TEs, iv) diploid genes without nearby TEs.

Focusing on diploid genes, the pattern was consistent and clear: among the group of genes without TEs, a higher percentage were expressed (Fig. 4A) and expressed at higher levels (Fig. 4B) than genes with nearby TEs. This observation held across the six taxa and across individual RNA-seq samples (*SI Appendix,* Fig. S11 and S12). The difference could be striking; for example, in one leaf sample of Pinot Noir, 93.3% of diploid genes without a nearby TE were expressed, while only 71.4% of diploid genes with a nearby TE were expressed. The pattern in diploid genes was consistent with the findings that host silencing of TEs near genes often negatively affects expression of a neighboring gene (38) – e.g., siRNA-targeted TEs are associated with reduced gene expression (39) --and TEs close to genes may disrupt cis-regulatory elements such as enhancers and silencers that affect gene expression (40, 41).

**Fig. 4.**
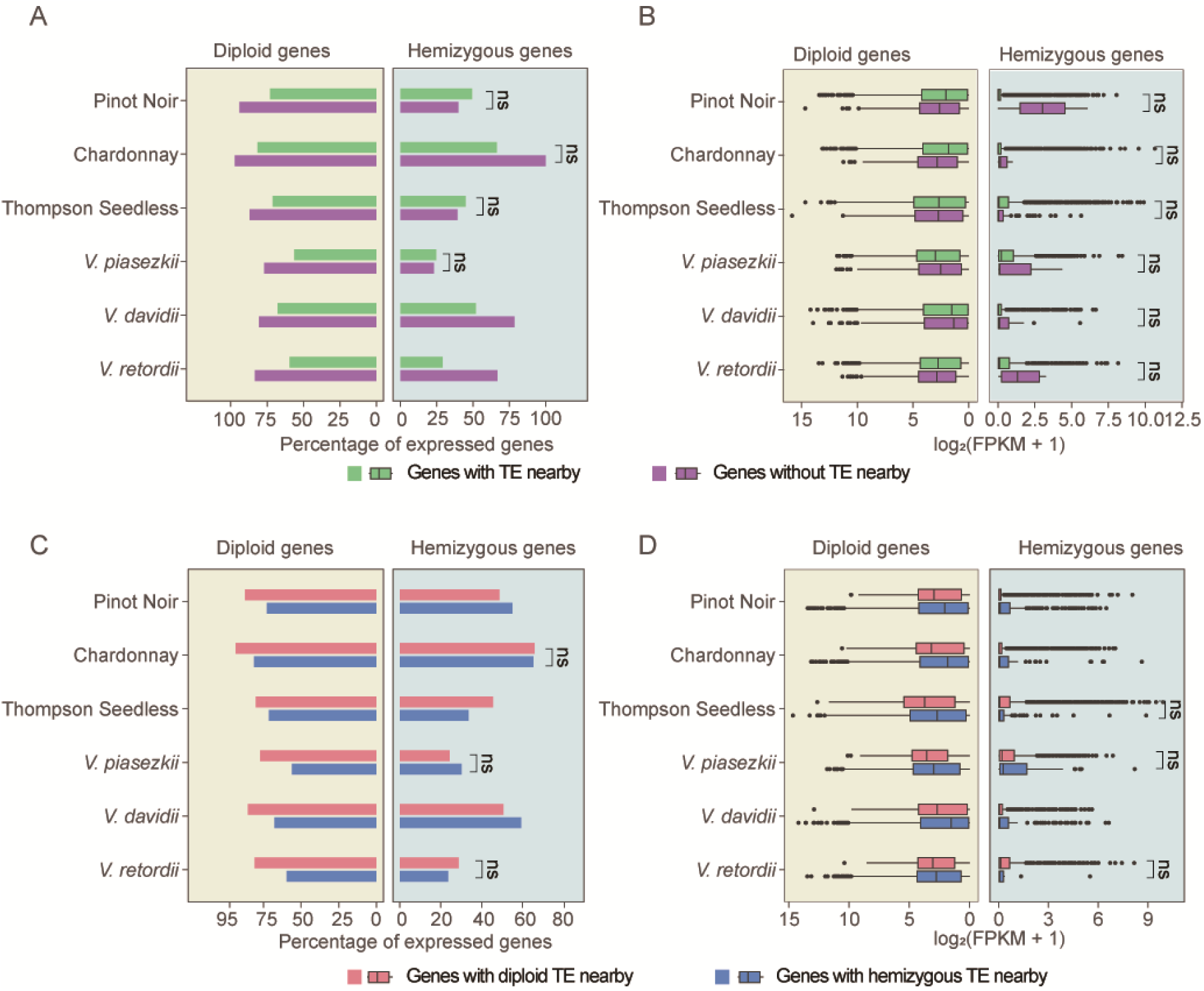
The cis-regulation of TE on hemizygous and diploid genes for clonal and outcrossing *Vitis* accessions. (A) The proportion of expressed hemizygous and diploid genes with or without nearby TEs. (B) Expression levels of expressed hemizygous and diploid genes with or without nearby TEs, shown as log_2_(FPKM+1). (C) The proportion of expressed hemizygous and diploid genes with nearby hemizygous or diploid TEs. (D) Expression levels of hemizygous and diploid genes with nearby hemizygous or diploid TEs, shown as log_2_(FPKM+1). A Fisher’s exact test was conducted for (A) and (C), while a Wilcoxon rank-sum test was applied to (B) and (D). Non-significant differences are indicated by "ns" (*P* > 0.05), while significant differences (*P* < 0.05) are shown without a mark. The line in (B) and (D) in the middle of the box represents the median, the edges of the box indicate the first and 3rd quartile, and the whiskers represent the range.

However, these patterns were less pronounced forhemizygous genes (Fig 4A, *SI Appendix,* Fig. S11 and S12). For examples, hemizygous genes near TEs tended to express more often in two taxa, *V. davidii* and *V. retordii* (Fig. 4A, Wilcoxon rank-sum test, *P* < 0.05), while the other four taxa showed no significant differences (Fig. 4A, Wilcoxon rank-sum test, *P* > 0.05). Meanwhile, there were no significant differences in the expression levels of hemizygous genes near TEs compared to those without nearby TEs in the six taxa (Fig. 4B, Wilcoxon rank-sum test, *P* > 0.05). Thus, the relationship between hemizygous genes and the presence of nearby TEs is less evident compared to that of diploid genes

Like genes, TEs can be diploid or hemizygous, so we also explored this effect on gene expression. To do so, we classified genes with nearby TEs into four categories: i) hemizygous genes with hemizygous TEs, ii) diploid genes with hemizygous TEs, iii) hemizygous genes with diploid TEs, and iv) diploid genes with diploid TEs. Across six taxa, the percentage of expressed diploid genes with nearby diploid TEs was generally higher compared to diploid genes with nearby hemizygous TEs (Fig. 4C and *SI Appendix,* Fig. S13) and at higher levels (Fig. 4D; *SI Appendix,* Fig. S14). Although the pattern for hemizygous genes was less obvious (Fig. 4D; *SI Appendix,* Fig. S14), the results generally suggest that SVs near genes (i.e., those resulting in hemizygous TEs) tend to reduce the expression of diploid genes more than nearby diploid TEs.

### Higher DNA methylation in hemizygous genes and TEs

One potential explanation for the effect of hemizygous versus diploid TEs has to do with epigenetic patterns – i.e., if hemizygous TEs are more highly methylated, they may more effectively dampen gene expression. We thus investigated DNA methylation patterns. We began by surveying hemizygous vs. diploid genes from leaves of four *Vitis* samples (Pinot Noir, Chardonnay, *V. piasezkii*, and *V. retordii*; *SI Appendix,* Table S9). For hemizygous genes in Pinot Noir leaves, we detected average weighted genomic DNA methylation levels of 45.8%, 23.0%, and 2.3% in the CG, CHG and CHH sequence contexts, respectively (Fig. 5A-5B and *SI Appendix,* Fig. S15); Similar to previous reports (42, 43), genic methylation levels were lower than the genome-wide methylation levels. For hemizygous versus diploid genes, the average DNA methylation level was 52.2% vs. 40.8%, 24.6% vs. 14.8%, and 2.3% vs. 1.7% in the CG, CHG, and CHH contexts, respectively. These patterns were largely consistent across taxa, and they generally reflect higher methylation levels for hemizygous genes compared to diploid genes.

**Fig. 5.**
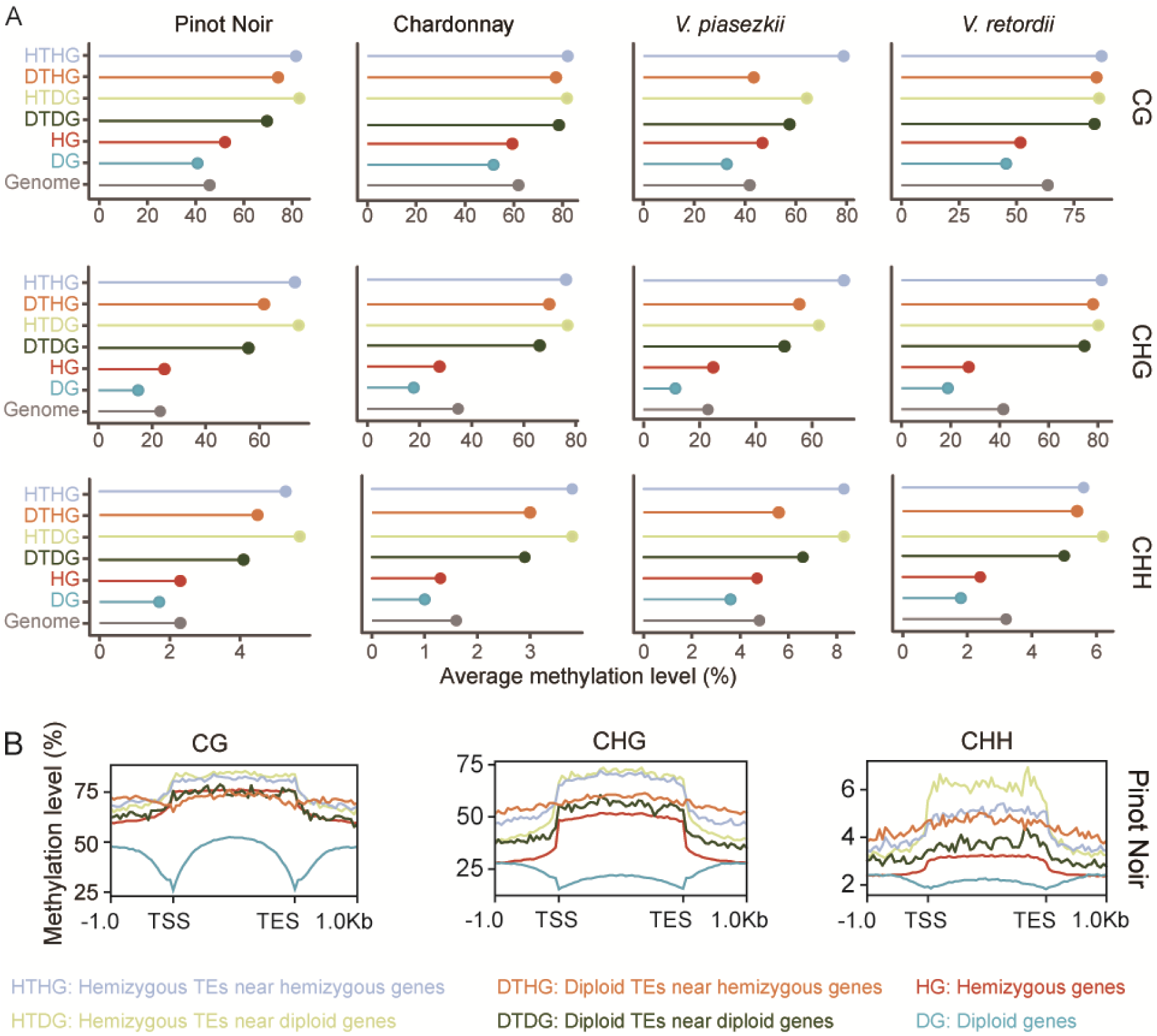
Epigenomic landscapes of hemizygous genes and their nearby diploid and hemizygous TEs. (A) Average DNA methylation levels across three methylation contexts (CG, CHG, or CHH) for different sequence types in four *Vitis* accessions. (B) Global distribution of DNA methylation levels for each sequence type for Pinot Noir. ‘TSS’ and ‘TES’ denote the transcription start and end sites of genes or the beginning or end of the TE annotations. The graphs include a 1-kb window upstream and downstream of each feature.

As expected, TEs were methylated at higher levels than genome-wide averages. However, it is interesting to note that hemizygous TEs close to diploid or hemizygous genes tended to be methylated at higher levels than diploid TEs close to diploid or hemizygous genes. For example, in the Pinot Noir sample, hemizygous and diploid TEs close to hemizygous genes have methylation levels of 81.6% and 74.2% in the CG context, 73.2% and 61.7% in the CHG context and 5.3% and 4.5% in the CHH context. Similar patterns were found in Chardonnay, *V. piasezkii*, and *V. retordii* (Fig. 5A). Hence, hemizygous TEs generally have higher methylation levels than diploid TEs, which may explain their stronger effect on nearby gene expression.

### Hemizygous gene expression levels correlated with gene body methylation

We then turned to the methylation status of individual genes. We defined gene body-methylated (gbM) genes as genes with CG methylation but without CHG and CHH methylation. We also categorized CHG methylated (mCHG) genes as genes with CHG methylation, and unmethylated (UM) genes as genes without CG, CHG and CHH methylation. Across two taxa, Chardonnay and *V. retordii*, the hemizygous genes tended to harbor a lower proportion of gbM genes than diploid genes. For instance, in Chardonnay, 18.1% of hemizygous genes (511 out of 2,821) and 28.8% of diploid genes (9,797 out of 33,980) were classified as gbM (Fig. 6A). In contrast, Pinot Noir and *V. piasezkii* exhibited a higher proportion of gbM genes in hemizygous genes compared to diploid genes. For example, in Pinot Noir, 26.7% of hemizygous genes (725 out of 2,711) and 21.8% of diploid genes (7,723 out of 35,459) were categorized as gbM (Fig. 6A). The difference between gbM proportions in hemizygous and diploid genes (i.e., hemizygous < diploid) and average CG genic methylation ratio pattern (hemizygous > diploid) in Chardonnay and *V. retordii* can potentially be attributed to the higher proportion of mCHG genes in hemizygous regions, which may influence their methylation status. For example, in Chardonnay, 54.4% of hemizygous genes (1,534 out of 2,821) and 23.5% of diploid genes (7,985 out of 33,980) were classified as mCHG genes, whereas in Pinot Noir, 42.3% of hemizygous genes (1,144 out of 2,711) and 17.3% of diploid genes (6,147 out of 35,459) were categorized as mCHG genes.

**Fig. 6.**
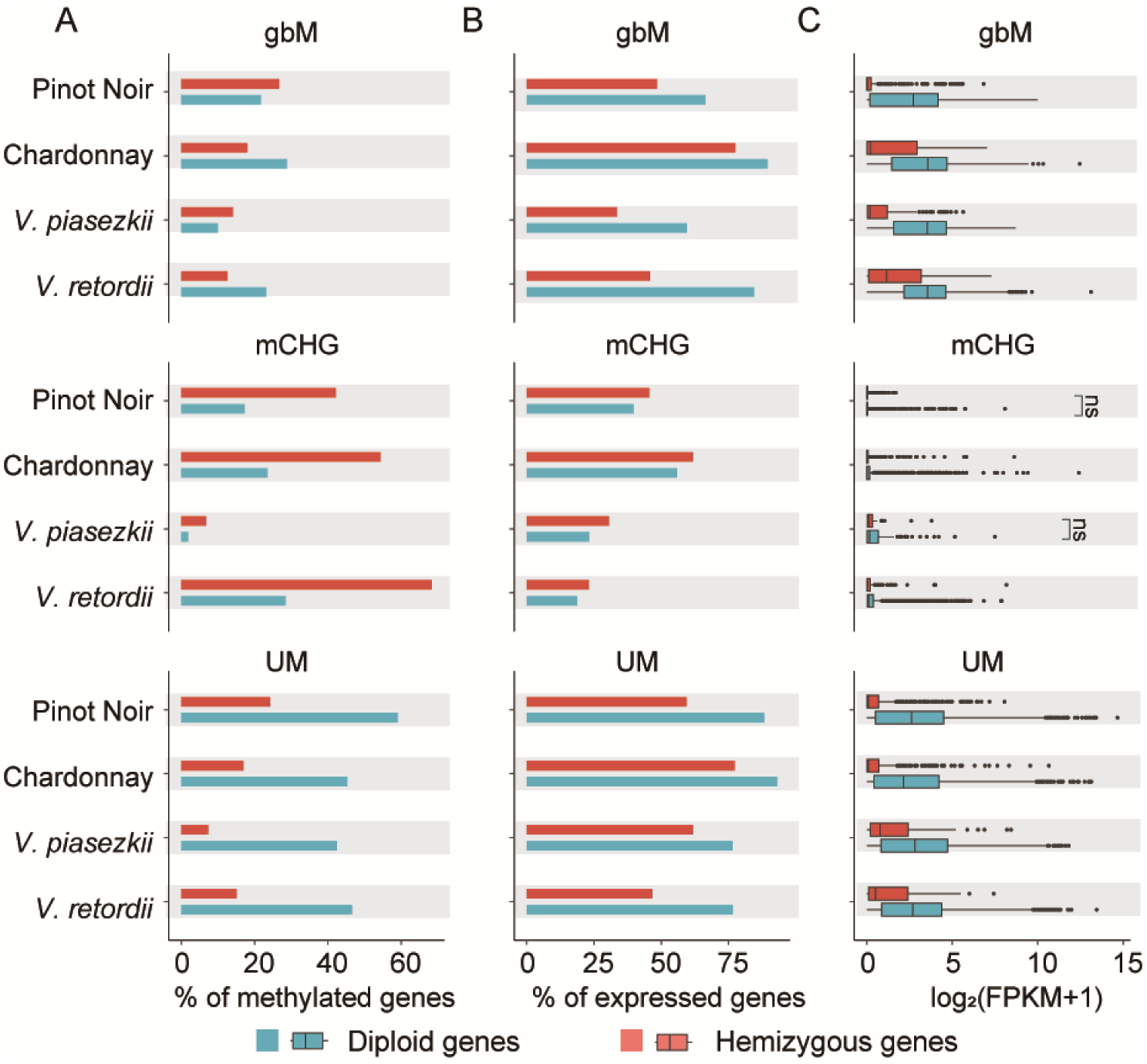
Epigenetic effects on the expression of hemizygous genes in four *Vitis* accessions. (A) The proportion of body-methylated genes (gbM), CHG methylated genes (mCHG) and unmethylated genes (UM) for hemizygous and diploid genes across four taxa. (B) The proportion of expressed hemizygous and diploid gbM, mCHG, and UM across four taxa. (C) Expression levels, shown as log_2_(FPKM+1), of expressed gbM, mCHG and UM. A Fisher’s exact test was conducted for (A) and (B), while a Wilcoxon rank-sum test was applied to (C). Non-significant differences are indicated by "ns" (*P* > 0.05), while significant differences (*P* < 0.05) are shown without a mark. The line in (C) in the middle of the box represents the median, the edges of the box indicate the first and 3rd quartile, and the whiskers represent the range.

After identifying gbM, mCHG, and UM genes, we investigated their expression patterns and observed several distinct trends. First, a smaller proportion of mCHG genes were expressed compared to gbM and UM genes (Fig. 6C), regardless of whether they were hemizygous or diploid. This finding aligns with previous studies indicating that mCHG methylation suppresses gene expression (42). The high proportion of hemizygous mCHG genes contributed to the overall lower expression levels of hemizygous vs. diploid genes (Fig. 3B). Second, a higher proportion of gbM and UM genes were expressed in diploid genes compared to hemizygous genes (Fig. 6B). Third, the patterns based on the proportion of expressed genes were largely reflected in expression levels. That is, mCHG genes were relatively lowly expressed, no matter if they were hemizygous or diploid (Fig. 6C); gbM and UM genes were expressed at higher levels than mCHG genes (Fig. 6C); and hemizygous gbM and UM genes were consistently expressed at lower levels than diploid genes (Fig. 6C).

## Discussion

Hemizygous genes have been studied extensively in sex-linked regions, but they can also occur beyond sex-linked regions of homologous chromosomes due to heterozygous SVs. Some SVs lead to the presence of a single allele on one homologous chromosome of an otherwise diploid organism. Here we have integrated genomic, transcriptomic and epigenomic analyses to estimate the frequency of these hemizygous genes, to explore their relationship to propagation history, and to characterize their features, expression, and epigenetic regulation.

### Hemizygous genes are most common in clonal lineages

Consistent with previous work, we have found that hemizygous genes are more common in clonal, as opposed to outcrossing lineages. Although hemizygosity has already been measured in a handful of plant taxa – i.e., primarily grape varieties and rice species – we have expanded observations to encompass nine clonally propagated cultivars from grapevine, apple, cassava, and potato, seven outcrossed samples from wild grapevine, wild apple, and wild rice, and four genomes expected to be fully homozygous (*SI Appendix*, Table S1). By focusing on hDELs, hBNDs, hDUPs, and hINVs relative to the reference assembly, we have documented, as expected, little evidence for hemizygosity in the homozygous samples, with estimates representing <1.2% of the genome (*SI Appendix*, Table S1). These results are not particularly surprising, but they show that we do not estimate high hemizygosity where there should be none.

In contrast to the homozygous samples, our work substantiates a growing consensus that outcrossing species can harbor a substantive portion of their genome as hemizygous. Among the seven outcrossed samples, 5.9%-11.7% of their genes are captured within hSVs, mimicking levels found in outcrossing rice and avocado. (Avocado is clonally propagated in cultivation, but the investigated tree had been produced by a recent outcrossing event.) In contrast, long-term clonal lineages consistently have an even more substantial fraction of their genomes and genes captured in a hemizygous state. Most of the observations to date have been based on grapevine clones, some of which have been propagated for 1000 or more years (44). However, by including cultivated apple, cassava, and potato, we have shown that this clonal phenomenon is not limited to grapevines (*SI Appendix*, Table S1). Moreover, the results accentuate how a traditional focus on inbred plants like *Arabidopsis thaliana*, rice, and tomato has biased our understanding of genetic variation. The inbred plants are typically highly homozygous with few sequence variants, but the genomes of clonal plants are highly heterozygous with genetic diversity that includes SVs and hemizygous genes (45).

High genetic variation in clonal lineages is not particularly unexpected, for two reasons. First, previous work on SVs has inferred, based on population samples, that they tend to be deleterious (22, 28). Second, forward simulations have consistently revealed that heterozygous, deleterious variants are expected to accumulate over time in clonal lineages, a phenomenon not observed in outcrossing plants (22, 46, 47). This accumulation reflects the fact that recessive deleterious alleles can hide as heterozygotes within a clonal lineage; whereas in outcrossing systems, they are expected to occasionally become homozygous and thus subject to selection. This accumulation also reflects that recombination is limited (i.e., effectively zero) in strictly clonal lineages, meaning that deleterious mutations cannot recombine onto different genetic backgrounds. Consistent with this argument, we find that hemizygous genes tend to be biased toward low-recombination, centromeric regions where selection is likely to be less efficient (Fig. 2A). Finally, it is interesting to note that previous work showed that domesticated, clonally propagated cassava has a marked 26% higher genomic burden of putatively deleterious nucleotides compared with its wild congener (48), consistent with its substantial burden of hemizygous genes (*SI Appendix*, Table S1).

Despite previous studies about the accumulation of deleterious variants in clonal lineages, the large number of hemizygous genes in clonal lineages is still somewhat surprising, because functionally hemizygous genes cannot (by definition) be recessive. Hence, the dynamics of the accumulation of hemizygous genes are likely to differ somewhat from the deleterious recessive case studied by forward simulation. Assuming that many (but not all; see below) of the SV events are slightly deleterious, several functional and evolutionary processes likely contribute to the accumulation of hemizygous genes in clonal lineages. One is a ratchet mechanism – i.e., once an SV occurs in a clonal lineage, it has only one possible fate, so long as it is not lethal, which is to remain in the clonal lineage. By this process, clonal lineages are expected to accumulate SVs. In theory, this accumulation is more likely when the SV events do not severely affect fitness; for that reason, we expect deleterious SVs to often have moderate functional effects.

### Hemizygous gene expression is moderated by epigenetic effects

Indeed, we have accrued evidence that hemizygous genes have moderate functional effects, based on three pieces of evidence. First, hemizygous genes are more likely to be non-expressed than diploid genes in our samples (Fig. 3A). That is, a higher proportion of hemizygous genes appear to be pseudogenes. Second, hemizygous genes are more likely to be members of gene families (Fig. 2E), implying that they are more likely to be functionally redundant. Thus, the loss of one copy of a multi-copy gene is likely to carry fewer fitness consequences than the loss of one allele of a critical single-copy gene. Finally, and somewhat surprisingly, as a group, hemizygous genes tend to be expressed at less than half the level of an average diploid genes, at about 20% (*SI Appendix*, Fig. S9C). This value is substantially less than the 50% expected of a single allele. It is hard to know the cause of this low expression pattern. It is possible, for example, that hemizygous genes are a biased sample that were lowly expressed in their diploid state before the SV event. Another possibility is that epigenetic effects act especially strongly on hemizygous genes to moderate their expression (see below).

In this context, it is worth accentuating that another set of hemizygous genes that have been studied intensively – i.e., sex-linked genes. Sex-linked genes tend to have an X:AA gene expression ratio of ∼0.5 in human, mouse, and nematode (11). Another possibility for sex-linked genes is dosage compensation, which predicts that hemizygous X-linked genes are expressed at twice the level of diploid genes per active allele to balance the gene dosage between the X chromosome and autosomes (12). The upregulation of the hemizygous copy may be sufficient to mitigate negative fitness effects, even if expression still falls significantly short of ancestral expression levels, and may also mitigate the effects of aneuploidy (49–51). In contrast, we do not see any overarching evidence of complete or even partial dosage compensation of hemizygous genes. Instead, the opposite is true: the expression of hemizygous alleles is substantially less expressed than the average diploid allele.

We suspect that lower expression is at least partially due to epigenetic phenomena, for three reasons. First, in all four samples investigated, for both diploid and hemizygous genes, nearby hemizygous TEs have elevated levels of DNA methylation relative to their nearby diploid TEs (Fig. 5A). Several phenomena may contribute to this observation, including that hemizygous TEs may be more recent insertions (and therefore more actively targeted by host epigenetic responses) (Fig. 2D). Whatever the cause, the data hint that hemizygous TEs differ quantitatively in their methylation effects. Second, hemizygous genes also exhibit higher levels of methylation than diploid genes, specifically a higher proportion of mCHG alleles (Fig. 6A), which are usually a mark of low expression (Fig. 6C). Finally, we have shown that genes near TEs are consistently more lowly expressed than genes far from TEs (Fig. 4B), but this effect is more prominent for genes near hemizygous TEs (Fig. 4D). This may be a partial explanation as to why genes close to SVs are associated with reduced gene expression levels in other species, like tomato (52).

These observations have interesting parallels to previous studies that have suggested that DNA methylation is correlated with reduced gene expression levels for sex-limited genes on the Y or W chromosome (53). High levels of DNA methylation have also been associated with sex chromosomes in sticklebacks and papaya (18, 19). In addition, DNA methylation is a key feature in X-chromosome inactivation (20). These results suggest some similar features of DNA methylation patterns between sex-linked and non-sex-linked hemizygous genes. For example, hemizygosity in human autosomes is linked to decreased gene expression and increased methylation, and these phenomena contribute to cancer and disease. For example, 45% to 60% of human gastric cancer cells show reduced *RUNX3* expression due to a hemizygous deletion and promoter hypermethylation (33). Similarly, promoter hypermethylation and a hemizygous deletion leads to *KLF4* down-regulation and apoptosis induction, enhancing its antitumor activity (34). Finally, a hemizygous deletion containing the *TP53* gene is found in over 10% of newly diagnosed multiple myeloma patients, and it is associated with decreased expression, impaired *p53* response, and resistance to apoptosis due to promoter hypermethylation (35). Clearly, we cannot be certain what, if any, epigenetic mechanisms might be shared between sex-linked and human disease hemizygosity and that which we have studied here, but it is an interesting question for further research.

### Are hemizygous genes merely functional remnants?

Given the evidence that hemizygous genes tend to be shorter than diploid genes (Fig. S6D-6E), expressed at lower levels (Fig. 3B), potentially subjected to lower levels of purifying selection (as measured by *Ka/Ks*; Fig. 2B), preferably distributed in centromeric regions representing low recombination regions, and more heavily methylated (Fig. 4A and 4B), it is tempting to conclude that hemizygous genes are typically pseudogenes. Are they merely functional remnants of previously functional genes? Might they also simply represent the dispensable component of the genome? which tend to be less expressed and more methylated (54). While the answer to this question is likely ‘yes’ for most hemizygous genes, there is some tantalizing evidence suggesting that the answer may often be ‘no’.

Evidence supporting functionality of some hemizygous genes comes in a few forms. For example, a reasonable proportion of hemizygous genes have gbM patterns of methylation (Fig. 6A). In both hemizygous and diploid genes, gbM genes exhibit significantly higher expression levels compared to mCHG genes (Fig. 6C). Moreover, several studies have detected a correlation between the presence of gbM and the enhancement of gene or allelic expression (42), while others have found evidence that it is subject to natural selection based on population genetic arguments. In short, although the functional role of gbM (if any) is debated (55), it typically is a mark deposited and maintained on active genes (42). The fact that some hemizygous genes bear this epigenetic mark superficially suggests that they can be easily dismissed as non-functional.

In addition, hemizygous genes (as a group) demonstrate patterns of tissue/treatment-specific expression that are similar to those of diploid genes. This pattern does not hold at the single gene level, but nonetheless up to 50% of hemizygous genes exhibit tissue/treatment-specific expression in Pinot Noir (*SI Appendix,* Fig. S10). Of course, tissue/treatment-specific expression patterns are not proof of function, but it does indicate that some hemizygous are induced under different environmental and developmental conditions. Finally, there are some consistent patterns of GO enrichment, particularly for responses to biotic and abiotic stresses (Fig. 2F and *SI Appendix,* Fig. S8). Again, GO enrichment is not proof of function, but this evidence combines to make it reasonable to hypothesize that not all hemizygous genes are functional ‘junk’. Of course, the mere act of uncovering a recessive allele can have important functional consequences; we invoke again the compelling case of hemizygosity and the white berry phenotype of grapes (22, 30).

## Materials and Methods

### Sample selection and genome assembly and annotation

We used genome assemblies based on PacBio sequencing data for 22 diploid plant samples (*SI Appendix*, Tables S1-S3). The details of these materials are provided in *SI Appendix*.

Among the 22 genome assemblies, the haplotype-resolved assemblies of Pinot Noir (PN_AGIS2_hap1 and PN_AGIS2_hap2), Chardonnay (CHT2T_AGIS1_hap1 and CHT2T_AGIS1_hap2), *V. piasezkii* (PIA_AGIS1_hap1 and PIA_AGIS1_hap2) were newly generated for this study (*SI Appendix,* Table S2-S3). The plant material was grown at Agriculture Genomics Institute at Shenzhen (AGIS), Chinese Academy of Agriculture Science (CAAS). DNA extraction and the construction of SMRTbell libraries followed ref.

(24). SMRTbell libraries were sequenced on the PacBio Sequel II platform in the CCS mode. DNA extraction and the preparation of ultra-long ONT libraries followed ref. (25) and sequenced on the ONT platform. For Hi-C library construction, chromatin was digested with the restriction enzyme Mbol using a previously described Hi-C library preparation protocol (56). The Hi-C libraries were sequenced on an Illumina HiSeq X Ten Platform. The details are provided in *SI Appendix*.

HiFi and Hi-C reads of the three *Vitis* accessions (Pinot Noir, PN_AGIS_02; Chardonnay, CH_AGIS_01; *V. piasezkii*) were collectively assembled into haplotigs using Hifiasm v0.19.8-r603 with default parameters. The remaining contig-level assemblies were then anchored and oriented to 19 chromosomes based on PN_T2T genome similarity (57) using RagTag v2.1.0 (58). Meanwhile, the Hi-C reads were filtered using fastp v0.23.4 (59) with default parameters. The filtered Hi-C reads were anchored on the haplotigs with Juicer v1.6 (60) and transformed into Hi-C contact map using 3D-DNA v190716 (61). The Hi-C contact map was visualized in Juicebox v2.17.00 (62) and adjusted manually to generate the final chromosome scaffolds. We aligned HiFi and ONT reads (with ONT reads only for Chardonnay) to the genome using Minimap2 (v2.24-r1122) (63). Subsequently, we manually filled the gaps by examining the alignment using Integrative Genomics Viewer (IGV) (v2.13.1) tool (64). The complete pipeline for genome assembly and gap filling is available on our lab GitHub@zhouyflab (see Code availability). Finally, haplotype-resolved genome assemblies were produced for four diploid grape varieties, with Chardonnay genome achieving a haplotype-resolved telomere-to-telomere level (CHT2T_AGIS1_hap1 and CHT2T_AGIS1_hap2, *SI Appendix,* Fig. S1).

Gene annotations were generated using the MAKER (65) and Liftoff pipelines (66) (*SI Appendix*, Fig. S2) for all seven *Vitis* samples, which included three newly assembled accessions and four previously published one (57, 67, 68). This strategy identified between 38,058 and 43,935 genes, with BUSCO scores up to 98.4% across seven *Vitis* crops (*SI Appendix,* Table S4). We performed TE annotation using RepeatModler/RepeatMasker (RM) (69) and EDTA pipelines (70). For RepeatModeler, we generated a non-redundant TE catalog based on all haplotypes. RepeatMasker was used to execute homolog annotation. RM pipeline identified TE sizes ranging from 321 to 375 Mb, occupying 66.9% to 71.3% of the genome. Based on EDTA for de novo annotation, which identified 187 to 280 Mb, covering 35.1% to 53.3% of the genome (*SI Appendix,* Table S5).

The remaining 19 genome assemblies were retrieved from public resources (*SI Appendix*, Table S2). The assembly results are provided in *SI Appendix*.

### Identification and characterization of hemizygous genes

To identify hemizygous genes, we gathered the raw long-read PacBio HiFi data for the 22 genome assemblies (*SI Appendix*, Table S2). We then remapped reads exceeding 10 kb to each genome assembly and called SVs using a Sniffles (v2.0.6) (71) pipeline. In our pipeline, the PacBio HiFi reads were mapped onto genome assemblies using Minimap2 (v2.24-r1122) (63) with the MD and map-hifi flag, and variant callings were performed using Sniffles (v2.0.6) (71). SV analysis outputs (VCF files) were filtered based on the following three steps: (1) we removed SVs that had ambiguous breakpoints (flag: IMPRECISE) and low-quality SVs that did not pass quality requirements of Sniffles (flag: UNRESOLVED); (2) we removed SV calls shorter than 50 bp; (3) we removed SVs with less than four supporting reads. Hemizygous regions were defined as deletion regions with 0/1 flags based on SV inferences, and genes that 80% overlapped hemizygous regions were defined as hemizygous genes. Genes were extracted from hemizygous regions of the genome with bedtools (v2.30.0) intersect with command “bedtools intersect -wo -a hemizygous_regions.bed -b gene.bed -F 0.8”. The remaining genes were termed diploid genes. To assess the effect of sequencing depth on SV and hemizygous genes detection, PacBio HiFi reads were subsampled to a depth of approximately 10×to 80×across samples, based on reads exceeding 10 kb, using seqkit (v2.2.0) (72). The effect of sequencing depth on SV and hemizygous genes detection was plotted for each taxon (*SI Appendix,* Fig. S4).

To further filter hemizygous regions, we aligned the two haplotypes in each of the six *Vitis* accessions (*SI Appendix*, Table S1). Whole-genome alignments were performed using Mummer (73). All alignment results between two allelic chromosome pairs were obtained using Nucmer (parameters: -c 500, -b 500, and -l 100) with the -maxmatch option (73). Then, the alignment results were filtered using delta-filter (parameters: delta-filter -m -i 90 -l 100) and the results were converted into a tab-separated file using the show-coords (parameters: show-coords -THrd) subprogram. The hemizygous genes identified through both the Sniffle pipeline and whole-genome alignments were defined as reliable hemizygous genes.

We then characterized sequence features and chromosome context, such as exon number, length, the proportion in centromere and telomere, synonymous mutation rate (*Ks*), and non-synonymous/synonymous mutation ratio (*Ka/Ks*) of hemizygous and diploid genes, the gene age based on phylostratigraphy analysis, the LTR insertion time, and the proportion of single copy genes of hemizygous and diploid genes. Gene length was calculated as the length between transcription start and end.

To investigate tandem repetitive sequences and centromeric regions, we used Tandem Repeats Finder (TRF, version 4.09) (74) to generate the statistics of the number of repeats and position information. These statistics were combined to annotate the centromeric repeats and telomere repeats. We examined the overlap with hemizygous and diploid genes after extending the regions 1Mb upstream and downstream of the centromeres and telomeres.

*Ka* and *Ks* values were estimated using MCScanX (75) pipeline (https://github.com/wyp1125/MCScanX) based on grapevine-*M. rotundifolia* genome sequence comparisons. We downloaded the genome fasta and gene annotation gff file of *M. rotundifolia* (http://www.grapegenomics.com/pages/Mrot/download.php), The corresponding coding and protein sequences were converted from fasta and gff file using gffread (v0.12.7) (76). The BLASTP (77) was performed using protein sequences (*E*-value < 1e^-10^, top 5 matches) to search all possible homologous gene pairs between each species pair. The output files based on BLASTP analysis were used as inputs for MCScanX, and pairwise *Ka*, *Ks* values of syntenic homologous genes were estimated using the Perl script “add_ka_and_ks_to_collinearity.pl” in the MCscanX package, which implemented the *Nei-Gojobori* algorithm (78).

We estimated the insertion times for diploid and hemizygous TEs using LTR identification from EDTA. LTRs were classified as diploid or hemizygous based on structural variation inferences. MEGACC (79) was employed with the commands “megacc.exe -a muscle_align_nucleotide.mao -d -o” and “megacc.exe -a distance_estimation_pairwise_nucleotide.mao -d -o” using the Kimura 2-parameter model and a mutation rate of 1 × 10⁻⁶ per copy per generation to calculate the K value. These calculations were based on the *K* value.

To estimate whether hemizygous genes are of more recent origin, we employed a phylostratigraphic procedure to determine the emergence time of each gene. Specifically, we queried all available proteins from protein-coding loci in each of the six *Vitis* genomes against the proteomes of 12 phylogenetically representative species (*SI Appendix*, Table S7 and Fig. S7) using BLASTP with default parameters and an *E*-value threshold of < 10^-3^. The species tree of 13 species including *Vitis* (*SI Appendix*, Fig. S7) was derived from the One Thousand Plant Transcriptomes Initiative (2019) (80) and one prior study (81). Based on the BLASTP outputs, we categorized all protein-coding genes into 12 phylostrata according to the consensus phylogeny. Genes that did not align were classified into a 13^th^ phylostratum. The age of each *Vitis* gene was determined by mapping it onto the consensus phylogeny using the most distant BLAST match above the significance threshold (36).

Using *M. rotundifolia* as outgroup, we determined single copy orthologous genes by performing OrthoFinder (37).

GO analysis was performed using David (82). We employed a *P* value < 0.05 to represent significantly enriched terms.

### Dissection of hemizygous gene expression patterns

To understand how hemizygous genes responded to fruit development, organ differentiation, and biotic and abiotic stress stimulus, we examined 168 RNA-seq samples from six grapevine and one apple, totaling 691 Gb (*SI Appendix*, Table S8). Among these, we generated three RNA-seq datasets from *V. piasezkii* leaves and three from *V. retordii* leaves. The remaining 162 samples were from public resources and categorized into samples based on fruit development, stress response, and organ differentiation based on their original experimental conditions. Detailed information and NCBI references can be found in Table S8 and *SI Appendix*.

Raw RNA-seq reads were trimmed by quality using Trimmomatic (v0.39) (83) with the options: LEADING:3 TRAILING:3 SLIIDINGWINDOW:10:20 MINLEN:36. High-quality reads were mapped onto the primary genome assemblies using HISAT2 (v.2.2.1) (84) with default parameters. Raw count for each gene was calculated based on FeatureCounts (2.0.1) (85) with the option: -p -B (paired-end reads, single-end reads without -B) -C -t transcript -g gene_id. Gene expression was quantified in normalized FPKM with a custom R script using the GenomicFeatures (86) package in R 4.1.0. In each tissue/treatment, gene expression was averaged over the biological replicates in each surveyed crop. Expressed genes were defined as those with FPKM > 0.

### Exploration of cis-regulatory effects of TEs on gene expression

Based on the identification of repeat sequences, we explored the cis-regulatory effects of TEs on gene expression. For this purpose, we first assigned each TE to its closest gene when it was within 2 kb (the distance to either 5’ or 3’ end of gene with >= 0 kb and <2 kb) using command “bedtools closest -wo -a gene.bed -b TE.bed”, and thus genes were separated in four classes: hemizygous genes with nearby TEs, hemizygous genes without nearby TEs, diploid genes with Nearby TEs, diploid genes without nearby TEs. We divided genes near TEs into four categories: hemizygous genes with either hemizygous or diploid TEs, and diploid genes with either hemizygous or diploid TEs.

### Unveiling DNA methylation patterns of hemizygous genes

Bisulfite-seq (BS-seq) for four samples were either generated for this study or downloaded from public sources (*SI Appendix,* Table S9). The Chardonnay clone chosen for BS-seq was FPS 04, a clone commonly grown in California and throughout the world. The reference plant is located at Foundation Plant Services, University of California. Samples of *V. piasezkii* and *V. retordii* were planted and collected in AGIS. DNA was isolated with the Qiagen DNeasy Plant Mini kit, and bisulfite libraries were prepared as previously described (43). Libraries were pooled and sequenced (150 bp paired-end) on the Illumina HiSeq2500. As a control for bisulfite conversion, lambda-DNA was spiked into each library preparation to measure the conversion rate of unmethylated cytosines (0.5% w/w). For publicly available datasets, BS-seq were retrieved from leaves in Pinot Noir (PRJNA381300) (87), Chardonnay (PRJNA691261) (88),

BSseq reads were trimmed for quality and adapter sequences using Trimmomatic (v0.39) with the options: LEADING:3 TRAILING:3 SLIIDINGWINDOW:10:20 MINLEN:36. Low quality reads and reads less than 36 bp were discarded. Bismark (v0.23.1) (89), in conjunction with bowtie2 (v 2.1.0) (90) with default parameters were used to align trimmed reads to the respective genome reference.

The number of methylated and unmethylated reads per cytosine was calculated using the bismark_methylation_extractor in Bismark (v0.23.1) (89). Methylated cytosines were identified using a binomial test incorporating the estimated rates of bisulfite conversion errors (P<0.01 after Benjamini-Yekutieli FDR correction) (91). False methylation rates (FMR) for each library were estimated for each taxon as one previous study performed (43), FMRs were estimated using lambda-DNA or chloroplast DNA using MethylExtract (92). A minimum coverage of two was required at each cytosine to determine methylation status. DNA methylation distribution plots were generated with deepTools (93).

We defined body-methylated genes following the strategy of ref. (55, 94). Briefly, we quantified the level of DNA methylation for each protein-coding region for each context-CG, CHG, CHH. The protein-coding region was defined as the annotated translation start to the termination codon. Taking the CG context as an example, *n*_CG_ was the number of cytosine residues at CG sites with ≥2 coverage in the gene of interest, *m*_CG_ was the number of methylated cytosine residues at CG sites for the same gene, and *p*_CG_ was the average proportion of methylated cytosine residues across all genes. Assuming a binomial probability distribution, the one-tailed *P* value for the departure of CG methylation levels from average genic proportion of DNA methylation was calculated as:

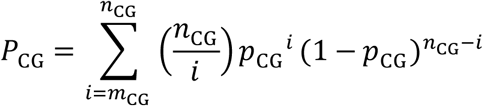

Where *P*_CG_ was a proxy of DNA methylation level. Using the same rationale, *P*_CHG_ and *P*_CHH_ were calculated for CHG and CHH context, respectively.

Given the binomial results, a gene was inferred to be gbM if CG methylation was significantly higher than the background (*P*_CG_ <=0.05), while CHG and CHH methylation were not significantly higher than the background (*P*_CHG_ > 0.05 and *P*_CHH_ > 0.05). Similarly, a gene was inferred to be CHG methylated if CHG methylation was higher than the background (*P*_CHG_ <= 0.05) and CHH methylation was not significantly higher than the background (*P*_CHH_ > 0.05). CHG methylated genes also tended to be CG methylated, but CG methylation was not required in our categorization. A gene was inferred to be CHH methylated if CHH methylation was higher than the background (*P*_CHH_ <= 0.05). CHH methylated genes also tend to be CG and CHG methylated. Finally, a gene was inferred to be unmethylated (UM) if CG, CHG, and CHH methylation were not significantly higher than the background (*P*_CG_ > 0.05, *P*_CHG_ > 0.05, and *P*_CHH_ > 0.05). In any other case, the gene methylation state was not inferred.

## Data availability

The PacBio CCS, ONT, Hi-C, RNAs-seq, BS-seq data have been deposited to the NCBI short reads achieved under the project number: PRJNA1178252 and the National Genomics Data Center (NGDC) Genome Sequence Archive (GSA) (https://ngdc.cncb.ac.cn/gsa/), with BioProject number PRJCA031656. The genome assembly and annotation have been deposited to zenodo: https://zenodo.org/records/14015567.

## Code availability

All scripts and codes performed in this study are available on GitHub: https://github.com/zhouyflab/Genomic_Epigenomic_Hemizygous_Crops.

## Acknowledgments

We thank R. Gaut for generating the BS-seq data and all members in the Zhou lab for the useful comments and discussions. This work was supported by the National Natural Science Foundation of China (No. 32372662), the Science Fund Program for Distinguished Young Scholars of the National Natural Science Foundation of China (Overseas) to Yongfeng Zhou, the National Key Research and Development Program of China (2023YFD2200700), Hainan Province Key Research and Development Project (ZDYF2024XDNY156), SZPU Research Project-6024330001K to Lin Tian, and the National Science Foundation grant (No.1741627) to Brandon S. Gaut.

## Author Contributions

Y.Z., B.S.G., and S.H. designed the research. Y.P., B.S.G., and Y.Z. wrote the manuscript. Y.P. performed the analyses. D.S, T.Z., X.S., and X.X. helped for genome assemblies and annotations. H.W., Y.W., L.C., Q.X., N.W., F.Z., Z.L., H.X., J.Y., L.T., B.S.G., Y.Z., W.H., S.C., and S.H. revised the paper. Y.L., X.F., Q.L., T.Z., X.X., and Y.P collected the data.

