## Supplementary material for "The genomic and epigenomic landscapes of hemizygous genes across crops with contrasting reproductive systems": SI text, Figure S1-S15, Table S1-S9

**This PDF file includes:**

Supporting text

Figures S1 to S15

Tables S1 to S9

Legends for Datasets S1 to S9

SI References

**Other supporting materials for this manuscript include the following:**

Datasets S1 to S9

### Supporting Information Text

### Supplementary Materials

#### Genome and HiFi Data resources

In this study, 22 samples with differing reproductive systems were utilized. Among them, the genomes of three accessions—**Pinot Noir** (PN_AGIS_02; *Vitis vinifera*), **Chardonnay** (CH_AGIS_01; *V. vinifera*), and ***Vitis piasezkii*** (PIA_AGIS_01)—were newly assembled. The remaining 19 sample accessions were retrieved from public databases, and the citations below refer to publications that produced the corresponding genome assemblies.

For four *Vitis* accessions—**Thompson Seedless** (*V. vinifera)*, ***V. retordii***, ***V. davidii,*** and **PN40024** (*V. vinifera*)—their genomes were assembled as described in references (1–4). For nine *Malus* accessions—**Golden Delicious** (*M. domestica*), **WA38** (*M. domestica*)**,** **Gala** (*M. domestica*), **M9** (*M. domestica*), **MM106** (*M. domestica*), **Fuji** (*M. domestica*), ***M. sieversii, M. sylvestris,*** and ***M. fusca***—their genomes were assembled as described in references (1-5). For one cassava accession—**TME204** (*Manihot esculenta*)—its genome was assembled as described in reference (6). For two *Solanum* accessions—**RH89-039-16** (*Solanum tuberosum*) and **SL5.0** (*S. lycopersicum* cv. Heinz 1706)—their genomes were assembled as described in references (7, 8). For three *Oryza* accessions—***O. rufipogon*** (Y476), **MH63** (*O. sativa*), and **Nipponbare** (*O. sativa*)—their genomes were assembled as described in references (9-11).

### Supplementary Method

#### The annotation of repeat sequences

To investigate tandem repetitive sequences and centromeric areas, we used Tandem Repeats Finder (TRF, v4.09) (12) to generate the statistics of the number of repeats and position information. These statistics were used to annotate the centromeric repeats and telomere repeats.

Transposable elements (TEs) were identified by the combined multiple programs. We generate the non-redundant TE catalogs using the RepeatModeler (v2.0.4) (13) based on all haplotypes. The RepeatMasker (v4.1.2, -e rmblast -lcambig -pa 2 -s) was used to execute homolog annotation. In addition, TEs were annotated using the Extensive de novo TE Annotator (EDTA) v2.0.0 including long terminal repeats (LTR), terminal inverted repeats (TIR), long interspersed nuclear element (LINE) repeats, Helitron-like DNA transposons, and so on (14).

### Gene annotation

We implemented a uniform annotation workflow to enhance the prediction of gene structure of all assemblies. We combined transcript evidence from RNA-seq datasets across diverse tissues/treatments. To improve the annotation process, we employed Hisat2 (v2.10.2) (15) to align RNA sequencing reads against the repeat-masked assemblies. The mapping information were extracted using the StringTie (v1.3.0) (16) program. We mainly relied on the custom scripts integrated Braker (v3.0.1) (https://github.com/Gaius-Augustus/BRAKER), PASA (v2.5.1) (17) and MAKER (v3.01.03) (18) programs for our genome annotations. We developed gene models and conducted subsequent searches utilizing AUGUSTUS (v3.4.0) (19). In addition, we removed genes residing within duplicated regions, had coding sequence lengths below 50, or lacked supporting evidence. We then checked the incomplete genes and alternative splicing events and filtered low confidence gene structures based on hidden Markov models and Pfam (20) (v1.6) database. Besides, gene annotations of six *Vitis* crops also performed based on PN_T2T genome and annotation file as reference (21) using Liftoff (v1.6.3) (22) with default parameters. Genes not detected by MAKER were added and the final annotation file was then renamed using a custom Python script. The PN_T2T annotation was updated based on the PN_AGIS02 genome and annotation in the current work using Liftoff. We incorporated gene annotations transferred by Liftoff that were not present in the previous annotation, with the help of a customized Python script combined with gff3_sort (https://github.com/NAL-i5K/GFF3toolkit/blob/master/docs/index.rst) and gffutil (http://daler.github.io/gffutils/).

### Supplementary Figures

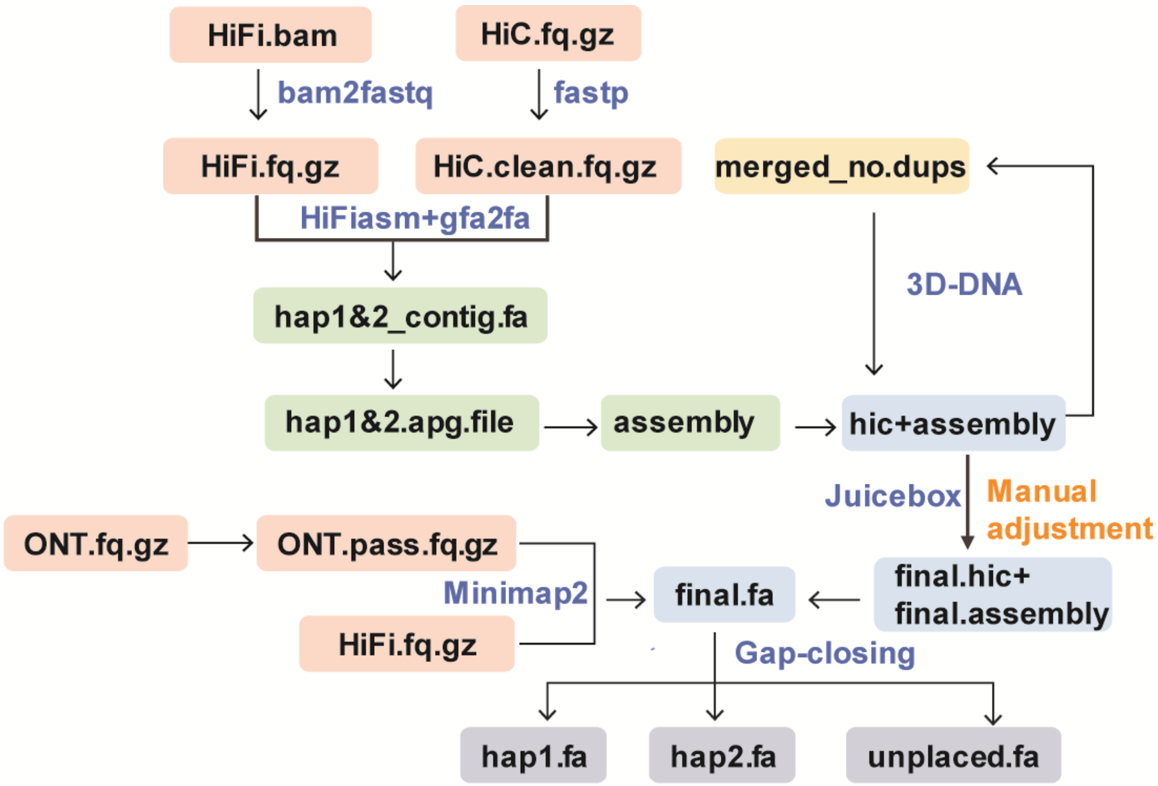

**Fig. S1.** Pipeline for Genome Assemblies. This figure illustrates the step-by-step process involved in assembling the genome.

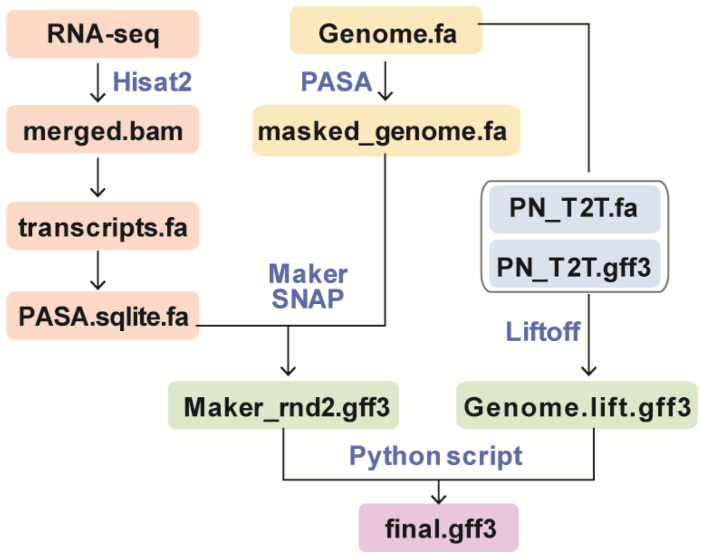

**Fig. S2.** Strategy for Genome Annotation of newly assembled genomes. In updating the PN_T2T genome annotation, we incorporated data from ref. (21) along with new annotations based on PN_AGIS2_hap1 using Liftoff.

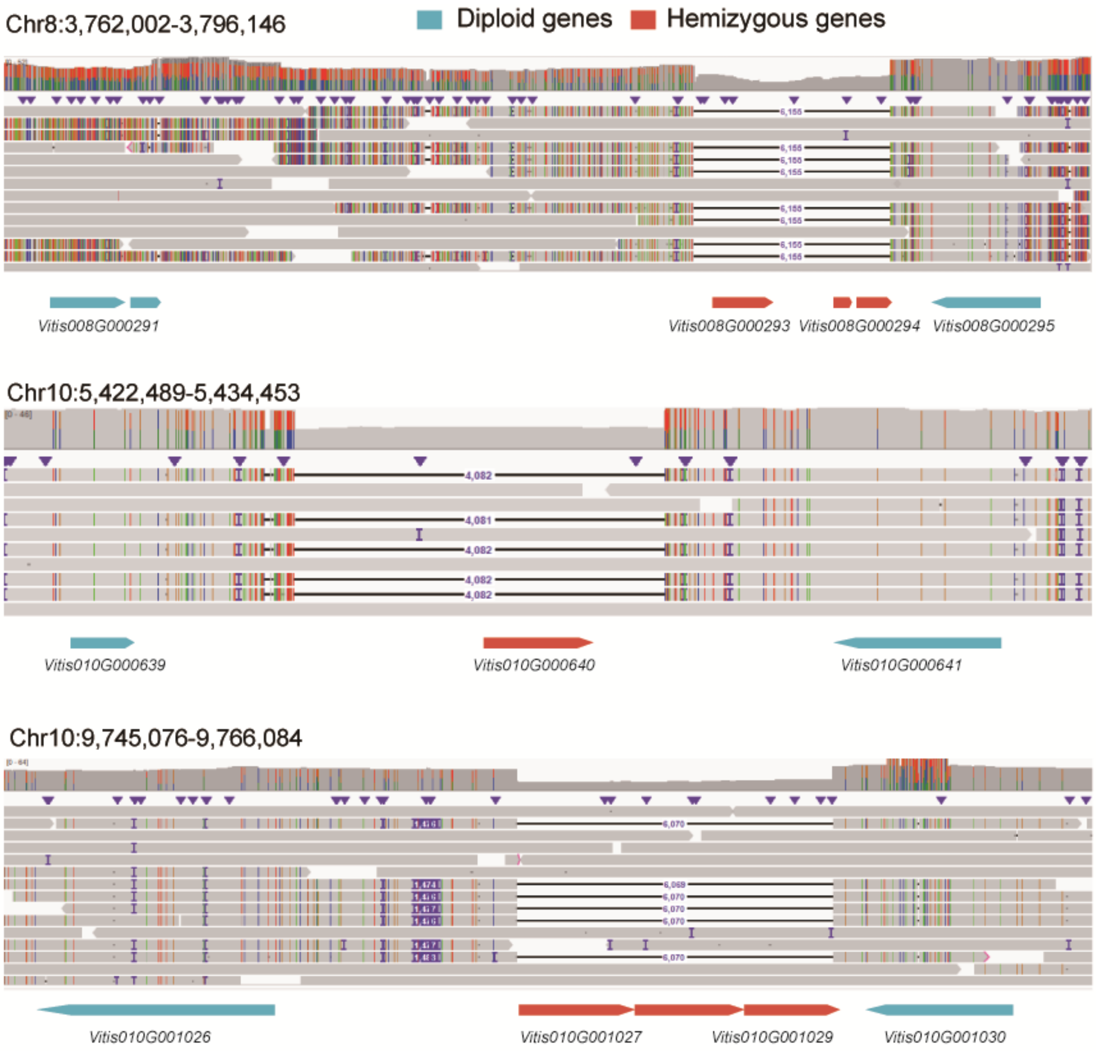

**Fig. S3**. IGV Examples of hemizygous and diploid genes in Pinot Noir.

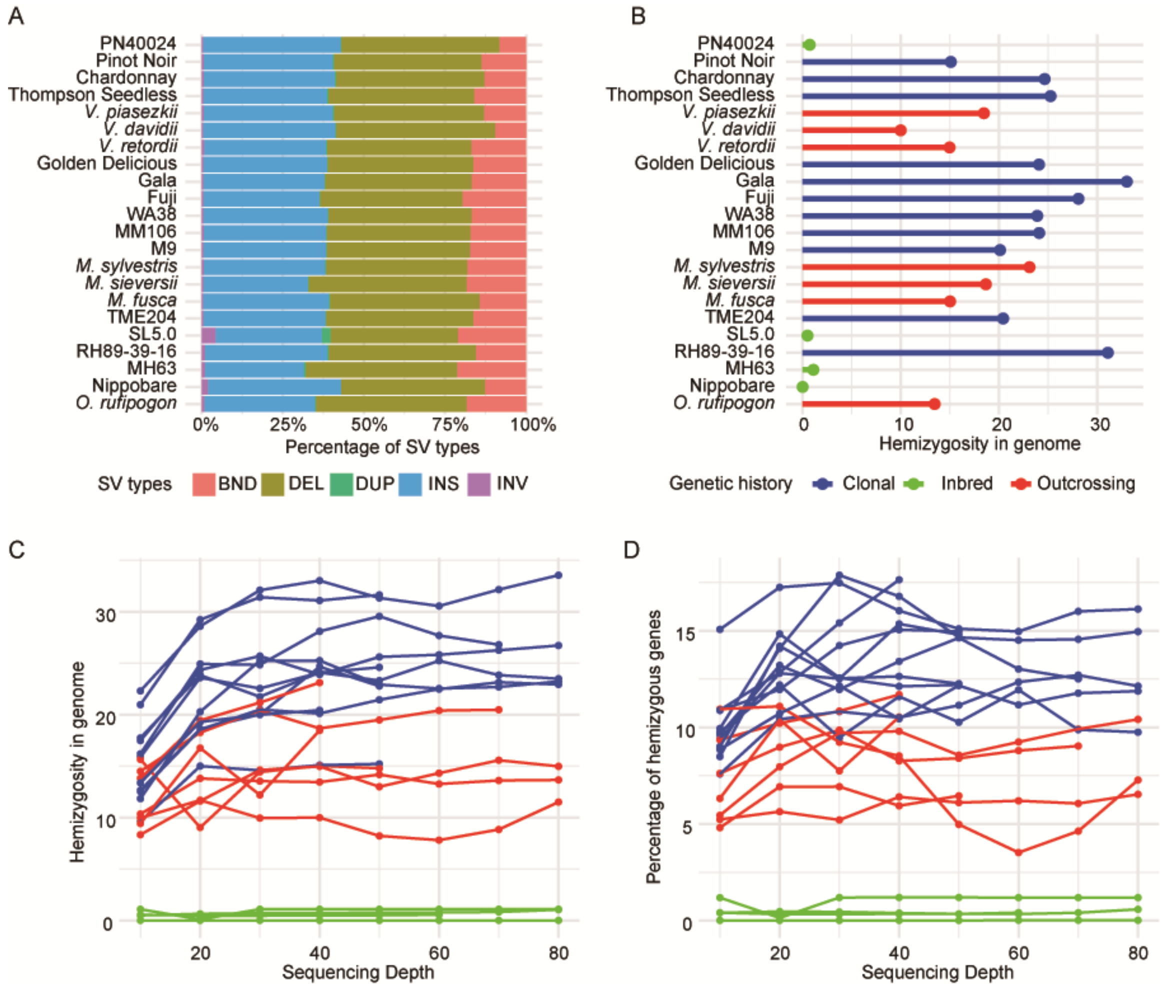

**Fig. S4.** The impact of sequencing depth. (A) Proportion of each heterozygous SV type within the total heterozygous SVs at a sequencing depth of 40×. (B) Estimation of genome hemizygosity at a sequencing depth of 40×. (C) Influence of sequencing depth using >10 kb HiFi data on genome hemizygosity, represented by the total size of heterozygous SVs relative to genome size. (D) Effect of sequencing depth on the proportion of hemizygous genes, inferred from SVs by mapping >10 kb HiFi data to the genome.

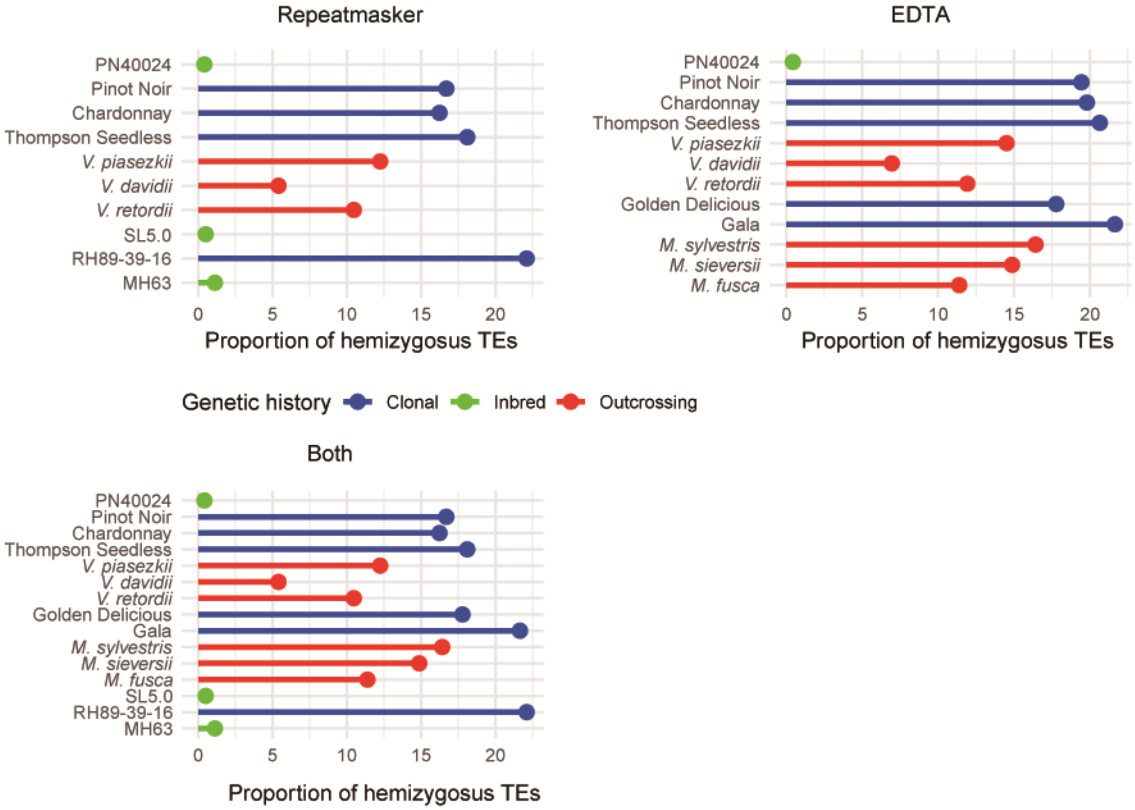

**Fig. S5.** The proportion of hemizygous TEs in genome based on RepeatMasker, EDTA and both methods.

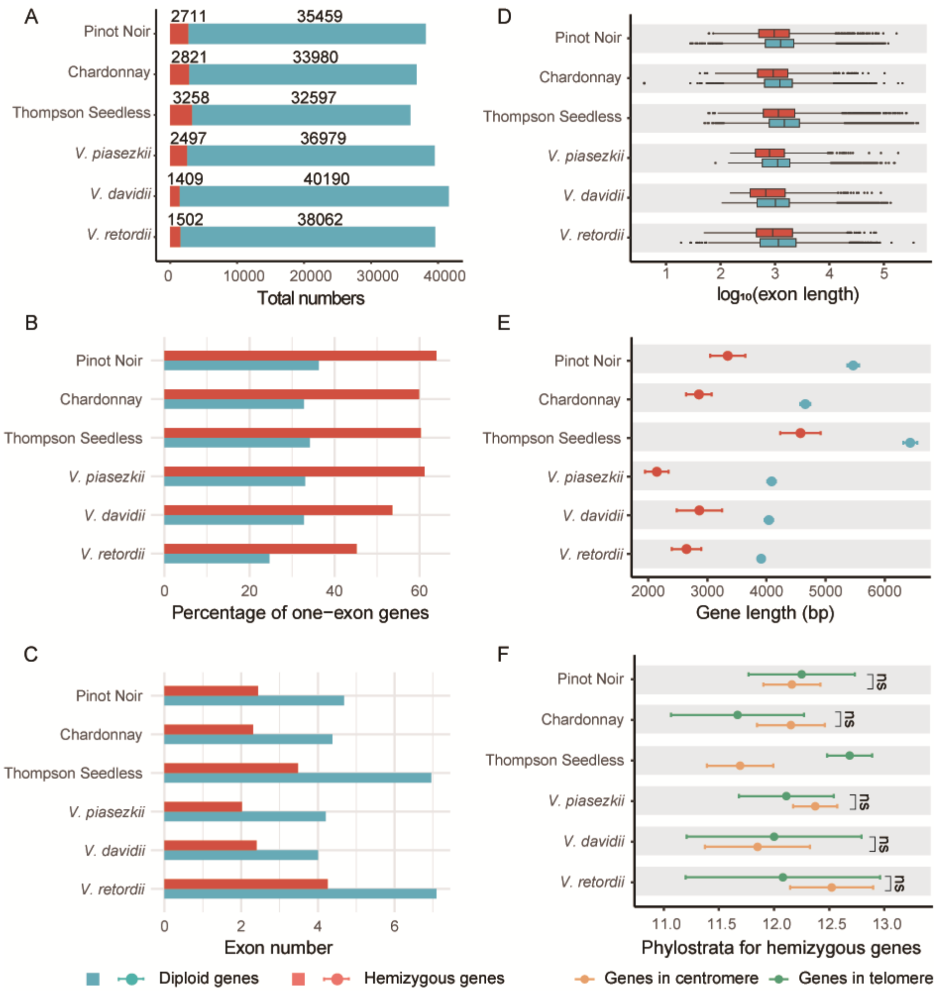

**Fig. S6.** Sequence Characterization of Hemizygous Genes. (A) Distribution of total hemizygous and diploid genes. (B) Percentage of hemizygous and diploid genes with a single exon. (C) Average exon number of hemizygous and diploid genes. (D) Exon lengths (bp), shown as log10(exon length), in hemizygous and diploid genes. (E) Average gene length (bp) in hemizygous and diploid genes. (F) Average phylostrata of hemizygous genes located in centromeric and telomeric regions. A Fisher's exact test was conducted for panel (A), while a Wilcoxon test was applied to panels (B–E). ns indicates $p > 0.05$, while significant differences are marked for other groups ($P < 0.05$). Error bars in panels (E) and (F) represent the 95% confidence intervals. In panel (D), the line in the middle of the box represents the median, the edges of the box correspond to the first and third quartiles, and the whiskers indicate the range of the data.

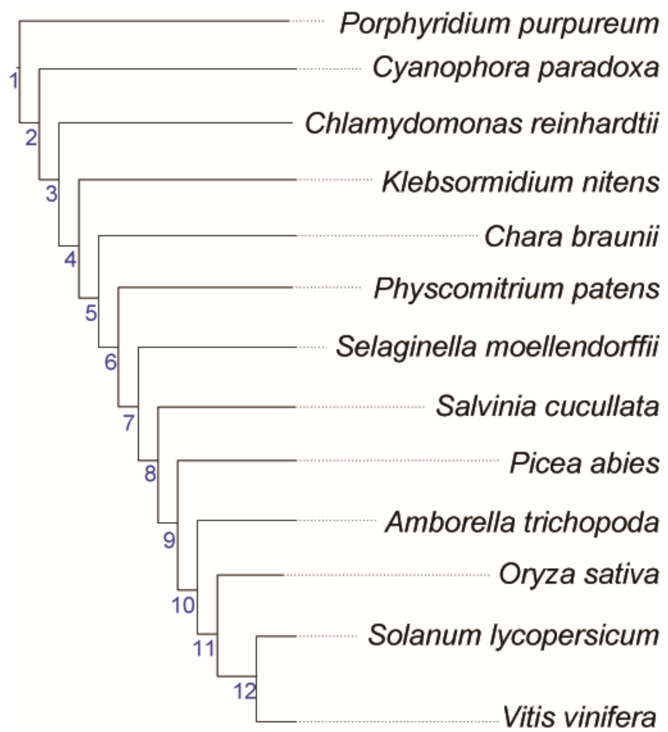

**Fig. S7**. Cladogram of the 13 species included in the analysis. The phylogenetic relationships are based on the One Thousand Plant Transcriptomes Initiative (2019). *V. retordii, V. piasezkii,* and *V. davidii* are in the sample place with *Vitis vinifera* in the species tree used for phylostratigraphic analysis. The numbers 1-12 in the phylogeny indicate the corresponding phylostrata.

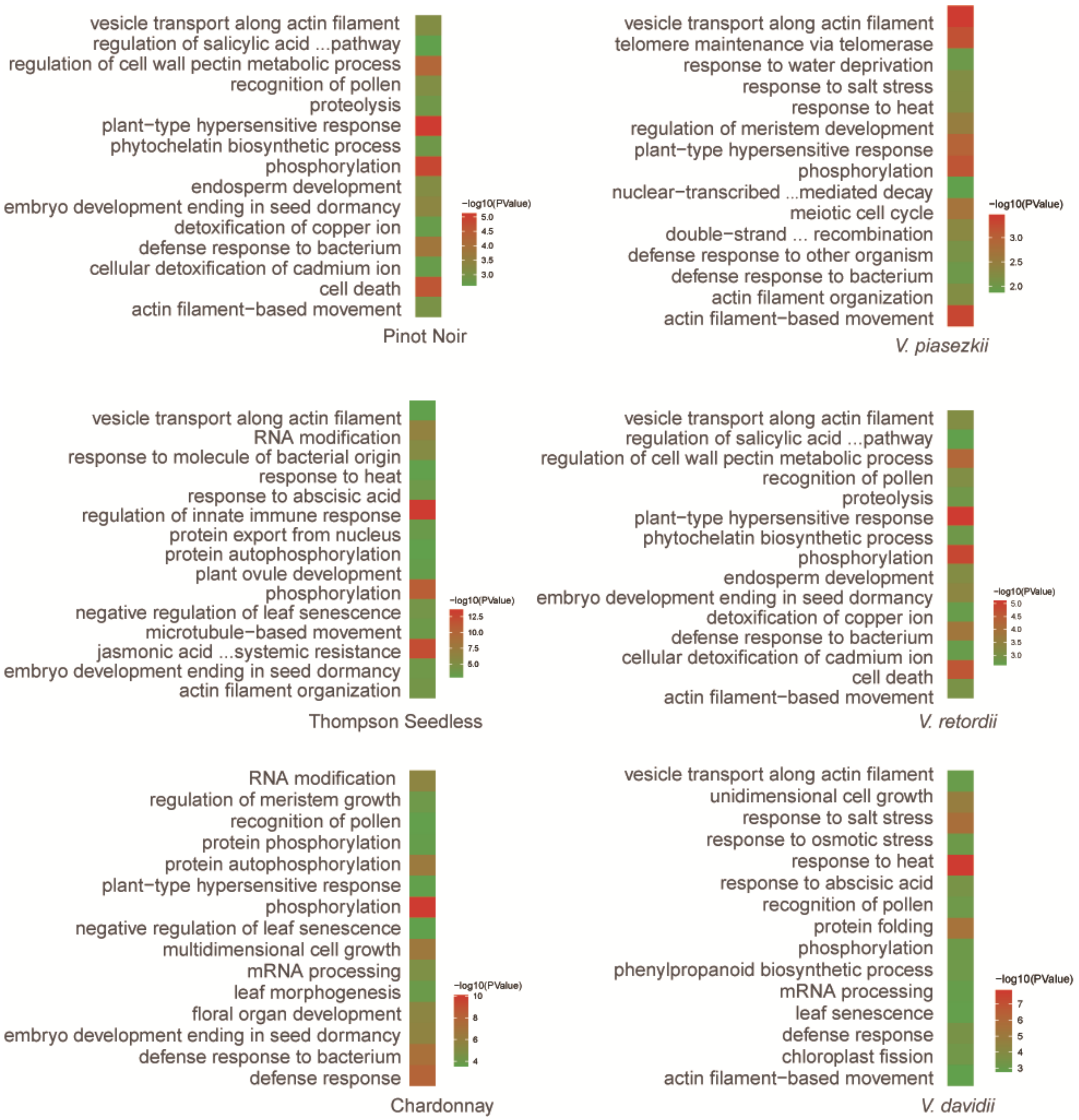

**Fig. S8.** The top 15 significantly enriched terms in biological processes for six *Vitis* samples.

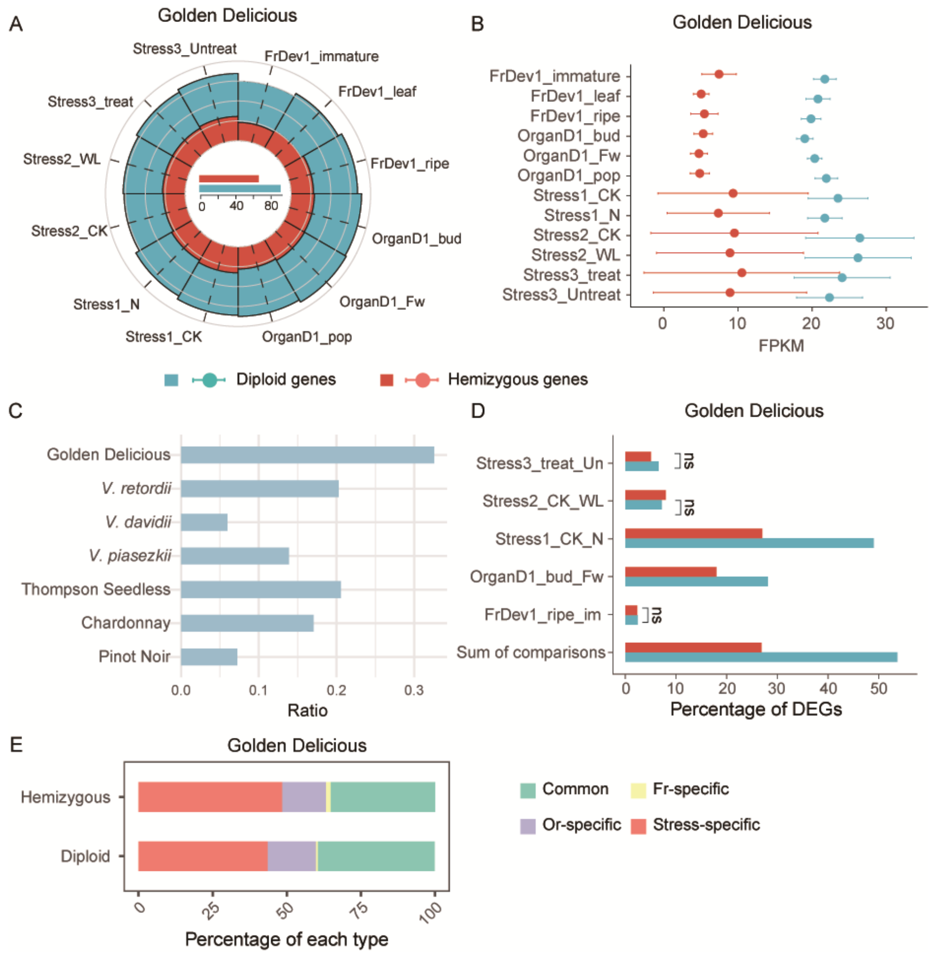

**Fig. S9.** Expression patterns of hemizygous genes. (A) The proportion of expressed hemizygous genes and diploid genes. Each circle represents 25% increments, the innermost is 25% and the outmost is 100%. (B) The expression level, shown as FPKM, for expressed hemizygous and diploid genes. Error bars show 95% bootstrap-based confidence intervals. (C) The distribution of hemizygous: diploid expression ratios. (D) The proportion of differentially expressed genes for hemizygous and diploid genes for Golden Delicious that allow control-treatment contrasts. (E) Proportion of common and unique differentially expressed hemizygous and diploid genes among three processes, including fruit development (Fr), organ differentiation (Or), abiotic and biotic stress stimulus processes (Stress). A Fisher test was conducted for A and D, and a Wilcoxon test was applied to B; ns: $p > 0.05$, while other groups showed significant differences ($p < 0.05$). Error bars indicated the 95% confidence interval.

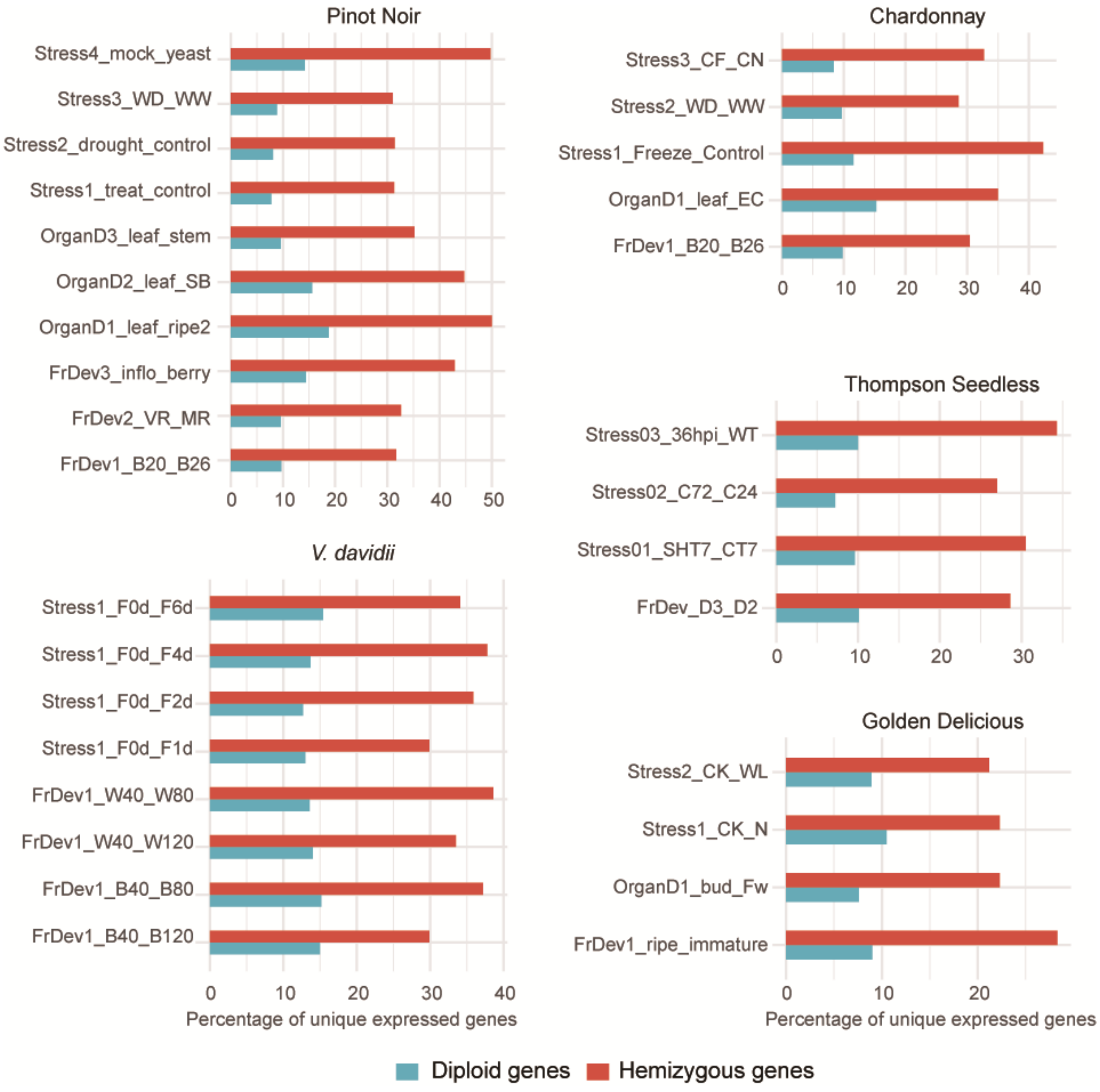

**Fig. S10.** The proportion of tissue/treatment-specific genes across five accessions. A Fisher's exact test was conducted, comparisons between hemizygous and diploid genes in all cases showed significant differences ($P$ < 0.05).

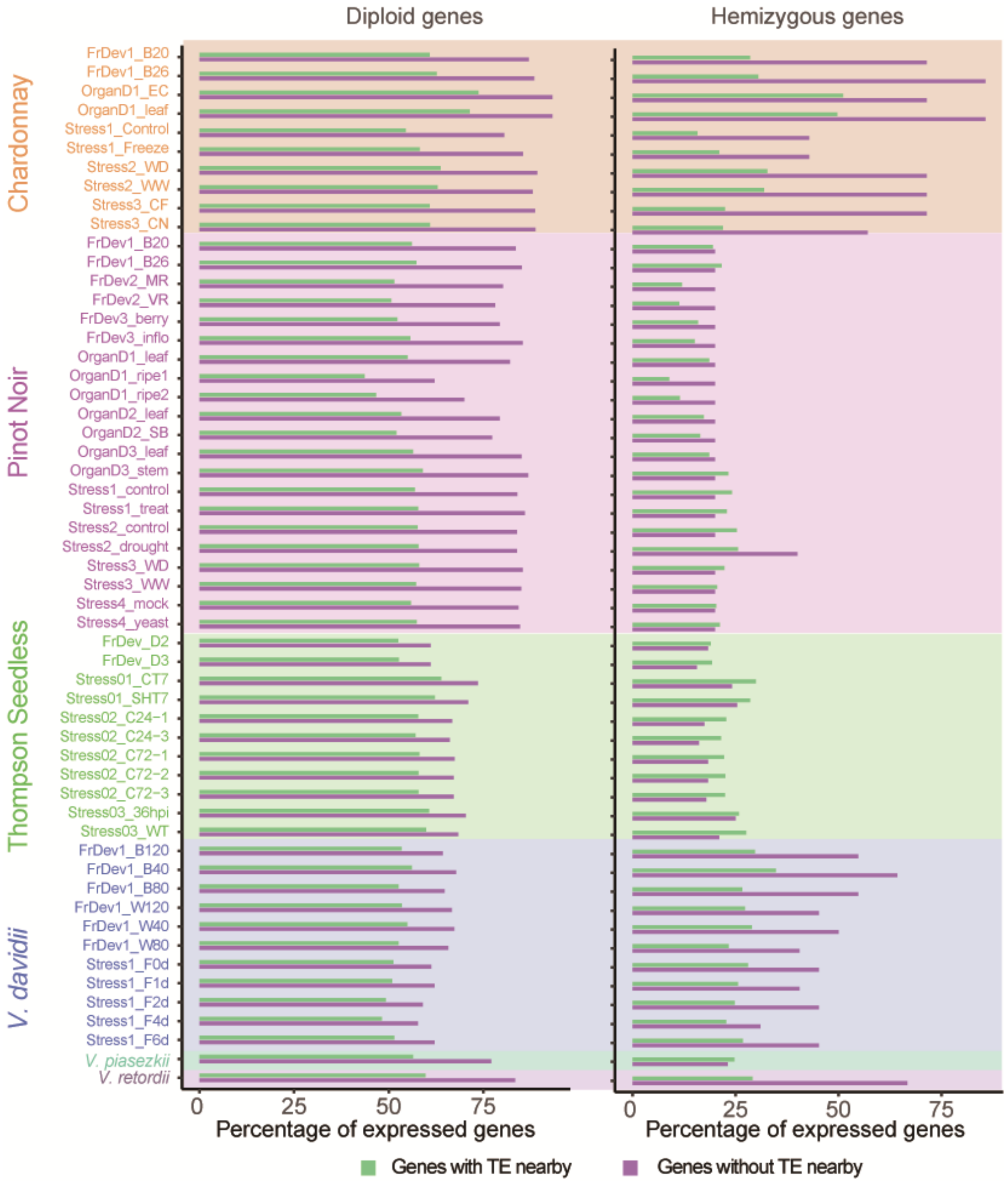

**Fig. S11**. The proportion of expressed proportion of hemizygous and diploid genes with nearby TEs and without nearby TEs compared to total hemizygous and diploid genes. A Fisher's exact test was conducted between genes with or without TE nearby. The *P* values are provided in Dataset S8.

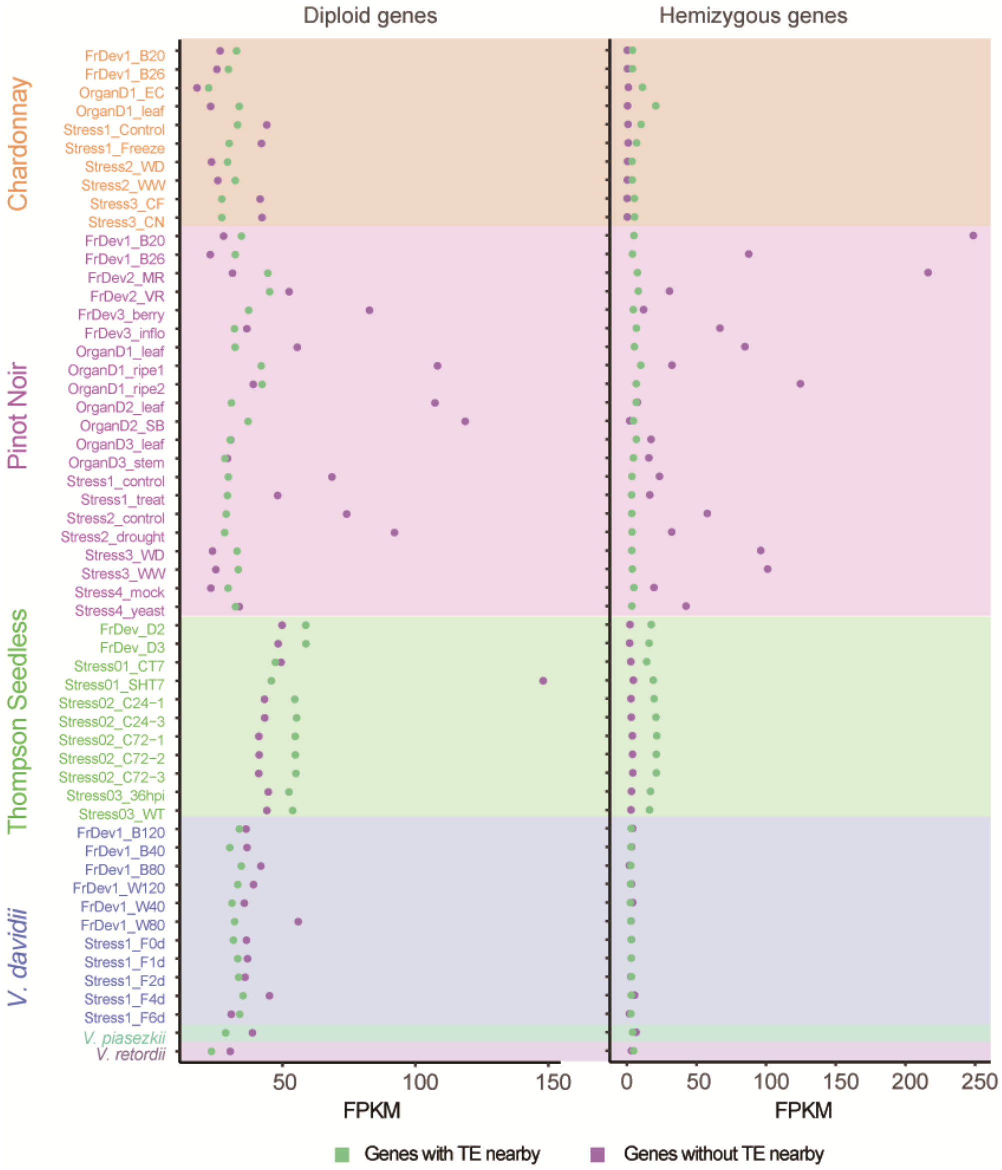

**Fig. S12.** The expression levels, show as FPKM, of hemizygous and diploid genes with nearby TEs and without nearby TEs. Error bars indicate the 95% confidence interval. A Wilcoxon rank-sum test was conducted between genes with or without TE nearby. The *P* values are provided in Dataset S8.

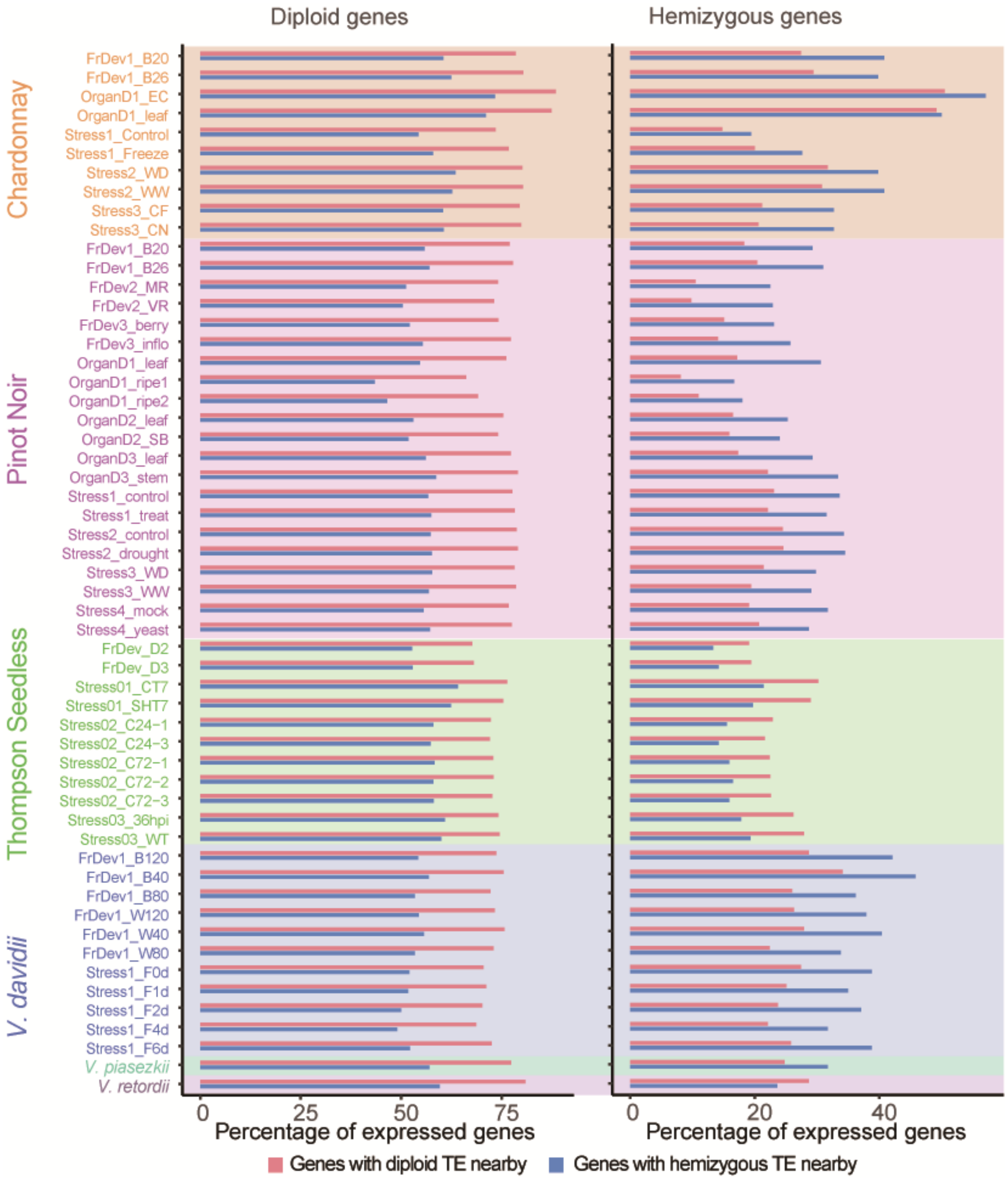

**Fig. S13.** The proportion of expressed of hemizygous and diploid genes with hemizygous and diploid TEs compared to total hemizygous and diploid genes. A Fisher's exact test was conducted between genes with or without TE nearby. The *P* values are provided in Dataset S8.

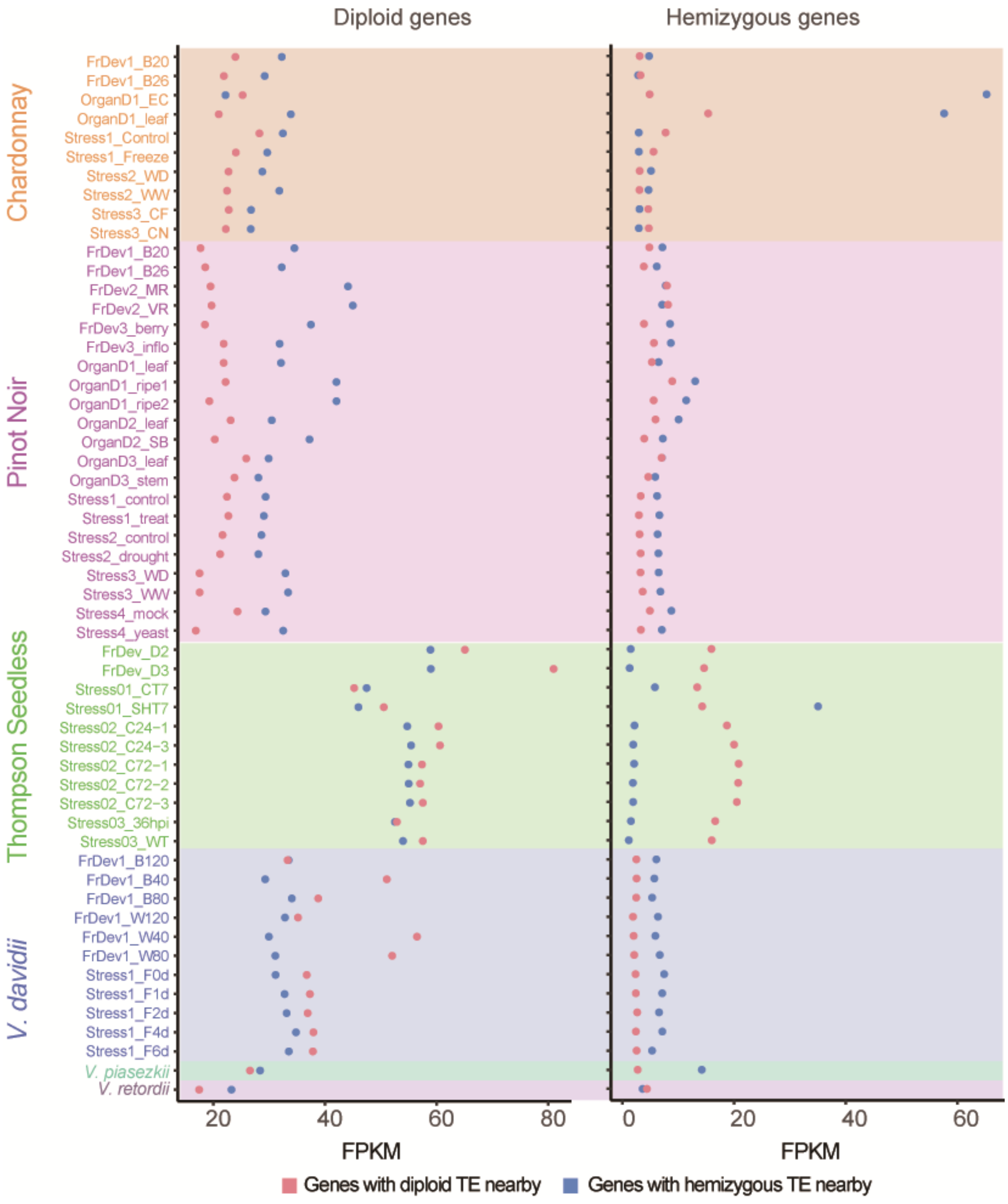

**Fig. S14.** The expression levels, shown as FPKM, of hemizygous and diploid genes with hemizygous and diploid TEs. Error bars indicate the 95% confidence interval. A Wilcoxon rank-sum test was conducted between genes with or without TE nearby. The *P* values are provided in Dataset S8.

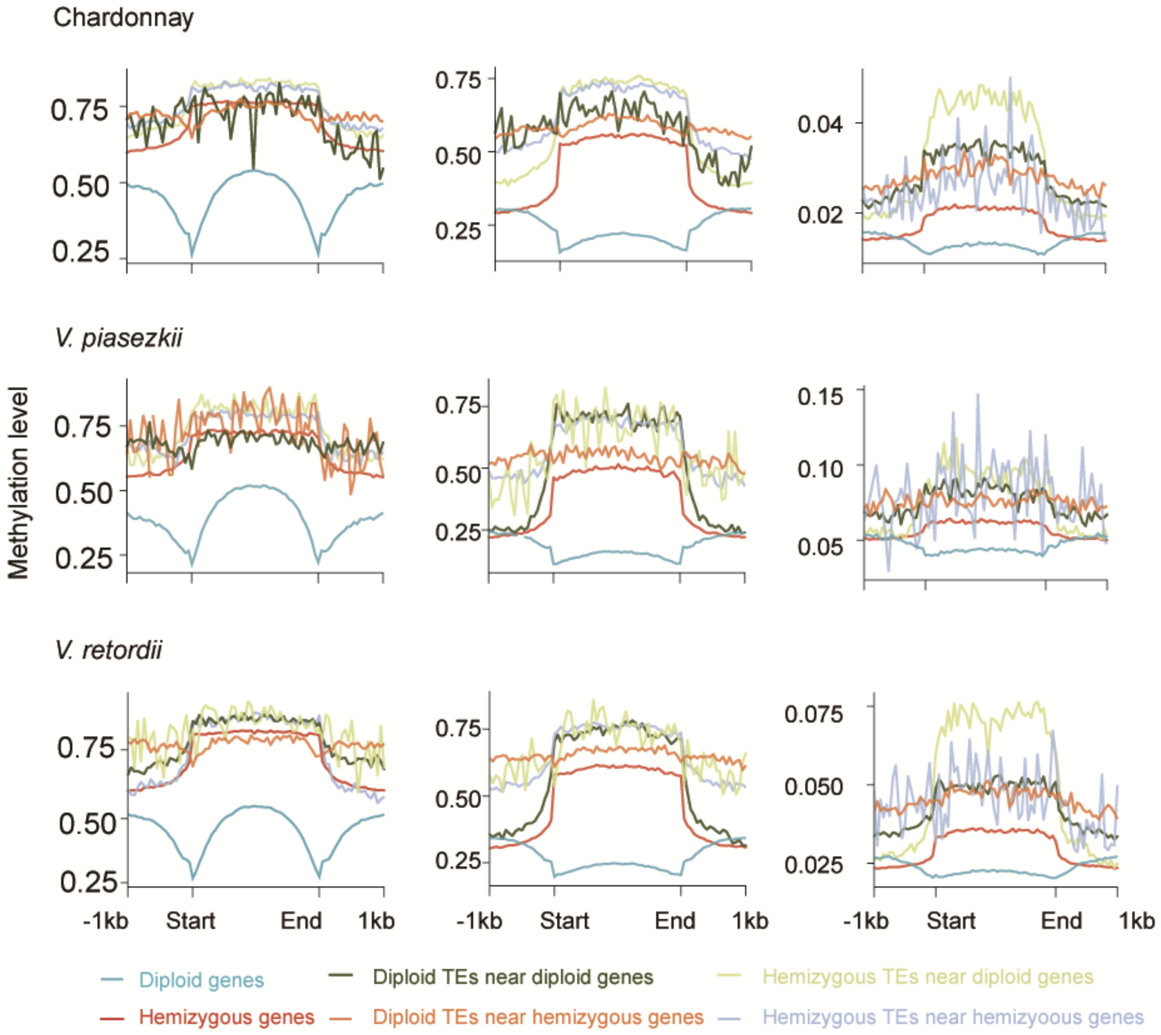

**Fig. S15**. Global distribution of DNA methylation levels in each sequence type. Start and end denote the transcription start and stop sites of genes or the beginning or end of the TE annotations.

**Table S1.** Summary of hemizygosity and percentage of hemizygous genes across different reproductive systems: eleven clonal, seven outcrossing, and four inbred.

| Reproductive system | Species | Accessions | Genome-wide Statistics | | | Hemizygous Genes | |
|---|---|---|---|---|---|---|---|
| | | | Size (MB)[3] | Hemizygous (Mb) | % | No. | % |
| Clonal | Grapevine (*Vitis vinifera*) | Pinot Noir | 493 | 76.0 | 15.4 | 5131 | 12.6 |
| | | Chardonnay | 500 | 124.6 | 24.9 | 6163 | 15.4 |
| | | Thompson Seedless | 485 | 124.4 | 25.6 | 6573 | 16.8 |
| | Apple (*Malus domesticata)* | Golden Delicious | 648 | 156.7 | 24.2 | 5380 | 12.1 |
| | | W38 | 648 | 154.1 | 23.8 | 5539 | 10.4 |
| | | Gala | 651 | 183.3 | 28.1 | 7403 | 16.0 |
| | | M9 | 617 | 217.2 | 35.2 | 5273 | 10.5 |
| | | MM106 | 651 | 155.0 | 23.8 | 5825 | 11.6 |
| | | Fuji | 651 | 131.1 | 20.1 | 6621 | 13.4 |
| | Potato (*Solanum tuberosum)* | RH89-39-16 | 805 | 251.9 | 31.3 | 5912 | 15.1 |
| | Cassava (*Manihot esculenta*) | TME204 | 734 | 155.8 | 21.2 | 5676 | 17.6 |
| Outcrossing | *V. piaseskii* | | 504 | 93.5 | 18.5 | 4356 | 10.5 |
| | *V. davidii* | | 525 | 52.5 | 10.0 | 3745 | 8.5 |
| | *V. retordii* | | 537 | 79.9 | 14.9 | 4136 | 9.8 |
| | *M. fusca* | | 635 | 95.8 | 15.1 | 2752 | 5.9 |
| | *M. sieversii* | | 633 | 122.0 | 19.3 | 3738 | 8.3 |
| | *M. sylvestris* | | 615 | 145.1 | 23.6 | 4974 | 11.7 |
| | *O. rufipogon* | | 495 | 55.3 | 11.2 | 2315 | 6.4 |
| Inbred | *V. vinifera* | PN40024 | 411 | 3.6 | 0.9 | 166 | 0.4 |
| | *O. sativa* | Nipponbare | 396 | 0.0 | 0.0 | 7 | 0.01 |
| | *O. sativa* | MH63 | 386 | 4.4 | 1.1 | 719 | 1.2 |
| | *S. lycopersicum* | Heinz 1706 (SL5.0) | 802 | 4.1 | 0.5 | 128 | 0.3 |

**Table S2**. Genome assembly and HiFi data resources. HiFi reads longer than 10 kb were utilized for hemizygous gene identification.

| Genus | Species | Accession | Reproductive system | BioProject | Bases (Gb) | Mean length (bp) | Bases (>10kb) (Gb) | Mean length (>10kb) (bp) |
|---|---|---|---|---|---|---|---|---|
| *Vitis* | *V. vinifera* | Pinot Noir | Clonal | PRJNA1178252 | 22.6 | 17678 | 22.5 | 17737 |
| *Vitis* | *V. vinifera* | Chardonnay | Clonal | PRJNA1178252 | 20.6 | 19044 | 20.6 | 19108 |
| *Vitis* | *V. vinifera* | Thompson Seedless | Clonal | PRJNA1021353 | 59.8 | 13744 | 59.4 | 13795 |
| *Vitis* | *V. piasezkii* | | Outcrossing | PRJNA1178252 | 20.6 | 17865 | 20.5 | 17892 |
| *Vitis* | *V. davidii* | | Outcrossing | CRA017609 | 55.0 | 16394 | 54.6 | 16578 |
| *Vitis* | *V. retordii* | | Outcrossing | PRJNA1082915 | 57.0 | 14710 | 56.5 | 14790 |
| *Vitis* | *V. vinifera* | PN40024 | Inbred | PRJNA882193 | 33.3 | 16187 | 33.3 | 16209 |
| *Malus* | *M. domestica* | Golden Delicious | Clonal | PRJNA983958 | 27.8 | 15871 | 27.7 | 15939 |
| *Malus* | *M. domestica* | WA38 | Clonal | PRJNA1072127 | 60.0 | 15501 | 58.8 | 15715 |
| *Malus* | *M. domestica* | Fuji | Clonal | PRJNA814760 | 53.2 | 11749 | 44.2 | 12448 |
| *Malus* | *M. domestica* | Gala | Clonal | PRJNA591623 | 57.8 | 12398 | 56.7 | 12542 |
| *Malus* | *M. domestica* | MM106 | Clonal | PRJNA814760 | 68.5 | 14780 | 65.6 | 15192 |
| *Malus* | *M. domestica* | M9 | Clonal | PRJNA814760 | 67.1 | 15448 | 66.2 | 15616 |
| *Malus* | *M. fusca* | | Outcrossing | PRJNA899490 | 21.7 | 15653 | 21.5 | 15740 |
| *Malus* | *M. sieversii* | | Outcrossing | PRJNA591623 | 43.1 | 11687 | 40.2 | 11942 |
| *Malus* | *M. sylvestris* | | Outcrossing | PRJNA591623 | 26.1 | 12854 | 24.4 | 13662 |
| *Manihot* | *M. esculenta* | TME204 | Clonal | PRJEB43673 | 31.3 | 20445 | 31.3 | 20457 |
| *Solanum* | *S. tuberosum* | RH89-39-16 | Clonal | PRJNA573826 | 30.9 | 13313 | 30.9 | 13332 |
| *Solanum* | *S. lycopersicum* | Heinz 1706 | Inbred | PRJNA733299 | 34.4 | 16613 | 34.4 | 16619 |
| *Oryza* | *O. rufipogon* | | Outcrossing | PRJNA1029807 | 11.4 | 15837 | 11.4 | 15850 |
| *Oryza* | *O. sativa* | MH63 | Inbred | PRJNA573706 | 25.3 | 21347 | 25.3 | 21362 |
| *Oryza* | *O. sativa* | Nipponbare | Inbred | PRJNA953663 | 33.0 | 18459 | 32.8 | 18709 |

**Table S3.** Genome assembly metrics.

| **Genus** | **Species** | **Accession** | **Reproductive system** | **Haplotype** | **Seqs** | **Size (Mb)** | **N50 (Mb)** | **Busco** |
|---|---|---|---|---|---|---|---|---|
| ***Vitis*** | *V. vinifera* | Pinot Noir | Clonal | hap1 | 19 | 497 | 26.3 | C:98.5%[S:97.1%,D:1.4%],F:1.0%,M:0.5%,n:1614 |
| | | | | hap2 | 19 | 490 | 25.2 | C:98.5%[S:97.1%,D:1.4%],F:1.0%,M:0.5%,n:1614 |
| | | Chardonnay | | hap1 | 19 | 505 | 26.8 | C:97.3%[S:95.8%,D:1.5%],F:1.2%,M:1.5%,n:1614 |
| | | | | hap2 | 19 | 495 | 25.1 | C:98.2%[S:96.5%,D:1.7%],F:1.1%,M:0.7%,n:1614 |
| | | Thompson Seedless | | hap1 | 19 | 493 | 25.1 | C:98.4%[S:96.8%,D:1.6%],F:0.9%,M:0.7%,n:1614 |
| | | | | hap2 | 19 | 478 | 25.1 | C:98.1%[S:96.8%,D:1.3%],F:1.1%,M:0.8%,n:1614 |
| | *V. piasezkii* | | Outcrossing | hap1 | 19 | 506 | 26.2 | C:98.4%[S:96.7%,D:1.7%],F:1.1%,M:0.5%,n:1614 |
| | | | | hap2 | 19 | 503 | 26.2 | C:98.0%[S:96.6%,D:1.4%],F:1.1%,M:0.9%,n:1614 |
| | *V. davidii* | | | hap1 | 19 | 525 | 28.1 | C:98.4%[S:96.8%,D:1.6%],F:1.0%,M:0.6%,n:1614 |
| | | | | hap2 | 19 | 526 | 28.3 | C:98.4%[S:96.3%,D:2.1%],F:1.0%,M:0.6%,n:1614 |
| | *V. retordii* | | | hap1 | 19 | 533 | 27.4 | C:98.6%[S:96.9%,D:1.7%],F:1.0%,M:0.4%,n:1614 |
| | | | | hap2 | 19 | 541 | 28.2 | C:98.7%[S:97.0%,D:1.7%],F:0.9%,M:0.4%,n:1614 |
| | *V. vinifera* | PN40024 | Inbreeding | primary | 19 | 495 | 26.9 | C:98.5%[S:97.1%,D:1.4%],F:1.0%,M:0.5%,n:1614 |
| ***Malus*** | *M. domestica* | Golden Delicious | Clonal | hap1 | 17 | 651 | 37.7 | C:99.0%[S:62.1%,D:36.9%],F:0.6%,M:0.4%,n:1614 |
| | | | | hap2 | 17 | 645 | 36.3 | C:98.9%[S:62.1%,D:36.8%],F:0.6%,M:0.5%,n:1614 |
| | | WA38 | | hap1 | 17 | 645 | 36.0 | C:99.0%[S:62.3%,D:36.7%],F:0.6%,M:0.4%,n:1614 |
| | | | | hap2 | 17 | 652 | 37.2 | C:99.0%[S:62.1%,D:36.9%],F:0.5%,M:0.5%,n:1614 |
| | | Fuji | | hap1 | 17 | 652 | 37.3 | C:99.1%[S:62.0%,D:37.1%],F:0.6%,M:0.3%,n:1614 |
| | | | | hap2 | 17 | 651 | 36.9 | C:99.0%[S:62.4%,D:36.6%],F:0.6%,M:0.4%,n:1614 |
| | | Gala | | hap1 | 17 | 658 | 38.0 | C:97.4%[S:62.4%,D:35.0%],F:0.7%,M:1.9%,n:1614 |
| | | | | hap2 | 17 | 577 | 36.4 | C:89.0%[S:62.8%,D:26.2%],F:1.5%,M:9.5%,n:1614 |
| | | MM106 | | hap1 | 17 | 643 | 38.2 | C:98.7%[S:62.3%,D:36.4%],F:0.6%,M:0.7%,n:1614 |
| | | | | hap2 | 17 | 659 | 37.8 | C:99.1%[S:62.3%,D:36.8%],F:0.5%,M:0.4%,n:1614 |
| | | M9 | | hap1 | 17 | 652 | 38.6 | C:99.0%[S:63.6%,D:35.4%],F:0.6%,M:0.4%,n:1614 |

| | | | | | | | | |
|---|---|---|---|---|---|---|---|---|
| | | | | hap2 | 17 | 651 | 36.7 | C:98.9%[S:62.9%,D:36.0%],F:0.5%,M:0.6%,n:1614 |
| | *M. fusca* | | Outcrossing | hap1 | 17 | 637 | 36.8 | C:98.8%[S:61.7%,D:37.1%],F:0.7%,M:0.5%,n:1614 |
| | | | | hap2 | 17 | 632 | 36.1 | C:98.9%[S:61.6%,D:37.3%],F:0.7%,M:0.4%,n:1614 |
| | *M. sieversii* | | | hap1 | 17 | 654 | 37.6 | C:97.2%[S:64.2%,D:33.0%],F:0.9%,M:1.9%,n:1614 |
| | | | | hap2 | 17 | 612 | 35.8 | C:93.0%[S:63.1%,D:29.9%],F:1.2%,M:5.8%,n:1614 |
| | *M. sylvestris* | | | hap1 | 17 | 628 | 36.2 | C:98.0%[S:63.1%,D:34.9%],F:0.7%,M:1.3%,n:1614 |
| | | | | hap2 | 17 | 601 | 35.2 | C:94.5%[S:65.9%,D:28.6%],F:1.8%,M:3.7%,n:1614 |
| ***Manihot*** | *M. esculenta* | TME204 | Clonal | hap1 | 18 | 762 | 35.5 | C:99.2%[S:92.3%,D:6.9%],F:0.4%,M:0.4%,n:1614 |
| | | | | hap2 | 18 | 706 | 34.2 | C:99.1%[S:93.2%,D:5.9%],F:0.4%,M:0.5%,n:1614 |
| ***Solanum*** | *S. tuberosum* | RH89-39-16 | Clonal | hap1 | 12 | 810 | 66.1 | C:90.8%[S:86.7%,D:4.1%],F:0.6%,M:8.6%,n:1614 |
| | | | | hap2 | 12 | 800 | 66.7 | C:90.7%[S:86.6%,D:4.1%],F:0.7%,M:8.6%,n:1614 |
| | *S. lycopersicum* | Heinz 1706 | Inbreeding | primary | 13 | 802 | 67.6 | C:98.6%[S:98.0%,D:0.6%],F:0.4%,M:1.0%,n:1614 |
| ***Oryza*** | *O. rufipogon* | | Outcrossing | hap1 | 12 | 411 | 33.5 | C:98.6%[S:96.5%,D:2.1%],F:0.9%,M:0.5%,n:1614 |
| | | | | hap2 | 12 | 412 | 33.9 | C:98.3%[S:96.3%,D:2.0%],F:1.0%,M:0.7%,n:1614 |
| | *O. sativa* | MH63 | Inbreeding | primary | 12 | 396 | 31.9 | C:98.7%[S:96.5%,D:2.2%],F:0.9%,M:0.4%,n:1614 |
| | | Nipponbare | Inbreeding | primary | 12 | 386 | 31.1 | C:98.7%[S:96.7%,D:2.0%],F:0.8%,M:0.5%,n:1614 |

**Table S4.** Metrics of *de novo* genome annotation for seven grapevine samples and one cassava sample, and number of centromeres and telomeres (Nce and Nte) identified in seven grapevine samples.

| Genus | Species | Accession | Reproductive system | Haplotyp e | Number | Busco | Nce | Nte |
|---|---|---|---|---|---|---|---|---|
| ***Vitis*** | *V. vinifera* | Pinot Noir | Clonal | hap1 | 40590 | C:97.4%[S:86.4%,D:11.0%],F:1.4%,M:1.2%,n:1614 | 17 | 37 |
| | | | | hap2 | 40715 | C:97.3%[S:86.3%,D:11.0%],F:1.4%,M:1.3%,n:1614 | 17 | 37 |
| | | Chardonnay | | hap1 | 40138 | C:95.0%[S:92.4%,D:2.6%],F:2.8%,M:2.2%,n:1614 | 19 | 38 |
| | | | | hap2 | 39481 | C:88.7%[S:86.5%,D:2.2%],F:8.0%,M:3.3%,n:1614 | 19 | 38 |
| | | Thompson Seedless | | hap1 | 39170 | C:98.4%[S:70.0%,D:28.4%],F:0.9%,M:0.7%,n:1614 | 19 | 35 |
| | | | | hap2 | 38058 | C:96.5%[S:68.8%,D:27.7%],F:1.1%,M:2.4%,n:1614 | 19 | 36 |
| | *V. piasezkii* | | Outcrossing | hap1 | 41335 | C:90.7%[S:88.8%,D:1.9%],F:5.6%,M:3.7%,n:1614 | 19 | 38 |
| | | | | hap2 | 40638 | C:89.8%[S:87.7%,D:2.1%],F:5.9%,M:4.3%,n:1614 | 19 | 38 |
| | *V. davidii* | | | hap1 | 43935 | C:90.3%[S:88.1%,D:2.2%],F:6.6%,M:3.1%,n:1614 | 19 | 38 |
| | | | | hap2 | 42216 | C:90.3%[S:88.0%,D:2.3%],F:6.6%,M:3.1%,n:1614 | 19 | 38 |
| | *V. retordii* | | | hap1 | 42198 | C:88.9%[S:60.2%,D:28.7%],F:7.9%,M:3.2%,n:1614 | 19 | 38 |
| | | | | hap2 | 40949 | C:89.6%[S:60.9%,D:28.7%],F:8.1%,M:2.3%,n:1614 | 19 | 38 |
| | *V. vinifera* | PN40024 | Inbred | primary | 41975 | C:98.0%[S:87.3%,D:10.7%],F:1.2%,M:0.8%,n:1614 | 19 | 36 |
| ***Manihot*** | *M. esculenta* | TME204 | Clonal | hap1 | 32183 | C:94.0%[S:84.2%,D:9.8%],F:3.9%,M:2.1%,n:1614 | | |
| | | | | hap2 | 32976 | C:95.0%[S:83.8%,D:11.2%],F:1.1%,M:3.9%,n:1614 | | |

**Table S5.** TE annotation metrics based on RepeatMasker and EDTA pipelines.

| Genus | Species | Accession | Reproductive System | Haplotype | Genome Sizes (Mb) | RepeatMasker | | | EDAT | | |
|---|---|---|---|---|---|---|---|---|---|---|---|
| | | | | | | **Numbers** | **Size (Mb)** | **Size %** | **Numbers** | **Size (Mb)** | **Size %** |
| *Vitis* | *V. vinifera* | Pinot Noir | Clonal | hap1 | 497 | 1058677 | 339 | 68.3 | 434229 | 243 | 49.0 |
| | | | | hap2 | 490 | 1054763 | 334 | 68.2 | 432289 | 239 | 48.9 |
| | | Chardonnay | | hap1 | 505 | 1076283 | 346 | 68.5 | 440782 | 249 | 49.2 |
| | | | | hap2 | 495 | 1076901 | 336 | 67.9 | 442690 | 258 | 52.1 |
| | | Thompson Seedless | | hap1 | 493 | 1074602 | 333 | 67.7 | 437519 | 225 | 45.8 |
| | | | | hap2 | 478 | 1046946 | 321 | 67.1 | 437525 | 225 | 47.1 |
| | *V. piasezkii* | | Outcrossing | hap1 | 506 | 1120767 | 342 | 67.7 | 492079 | 249 | 49.1 |
| | | | | hap2 | 503 | 1108122 | 340 | 67.6 | 474354 | 242 | 48.1 |
| | *V. davidii* | | | hap1 | 533 | 1127926 | 362 | 67.8 | 340718 | 187 | 35.1 |
| | | | | hap2 | 541 | 1133438 | 362 | 66.9 | 482185 | 267 | 49.4 |
| | *V. retordii* | | | hap1 | 525 | 880279 | 367 | 69.8 | 507822 | 271 | 51.7 |
| | | | | hap2 | 526 | 881170 | 375 | 71.3 | 507529 | 280 | 53.3 |
| | *V. vinifera* | PN40024 | Inbred | primary | 495 | 159834 | 328 | 66.4 | 468759 | 243 | 49.0 |
| *Malus* | | Golden Delicious | Clonal | hap1 | 651 | | | | 899128 | 402 | 61.7 |
| | | Gala | | hap1 | 658 | | | | 906175 | 324 | 49.3 |
| | *M. fusca* | | Outcrossing | hap1 | 637 | | | | 639775 | 352 | 55.3 |
| | *M. sieversii* | | | hap1 | 654 | | | | 966001 | 392 | 60.0 |
| | *M. sylvestris* | | | hap1 | 651 | | | | 910527 | 321 | 49.3 |
| *Solanum* | | RH89-39-16 | Clonal | hap1 | 810 | 1358797 | 562 | 69.3 | | | |
| | | Heinz 1706 | Inbred | primary | 802 | 1395143 | 554 | 69.0 | | | |
| *Oryza* | | MH63 | Inbred | primary | 396 | 616102 | 205 | 51.7 | | | 49.0 |

**Table S6.** Hemizygous TEs identified based on RepeatMasker and EDTA pipelines.

| Genus | Species | Accession | Reproductive system | RM | | EDTA | | Overlapped | | |
|---|---|---|---|---|---|---|---|---|---|---|
| | | | | **Hemizygous** | **%** | **Hemizygous** | **%** | **Number** | **Hemizygous** | **%** |
| *Vitis* | *V. vinifera* | Pinot Noir | Clonal | 176744 | 16.7 | 84409 | 19.4 | 429960 | 83522 | 19.4 |
| | | Chardonnay | | 174850 | 16.2 | 87268 | 19.8 | 438499 | 86370 | 19.7 |
| | | Tomphson Seedless | | 194536 | 18.1 | 90360 | 20.7 | 367895 | 82948 | 22.5 |
| | *V. piasezkii* | | Outcrossing | 137246 | 12.2 | 71390 | 14.5 | 486448 | 70444 | 14.5 |
| | *V. davidii* | | | 60949 | 5.4 | 23659 | 6.9 | 337284 | 23364 | 6.9 |
| | *V. retordii* | | | 92180 | 10.5 | 60521 | 11.9 | 502618 | 59846 | 11.9 |
| | *V. vinifera* | PN40024 | Inbred | 4828 | 0.4 | 1990 | 0.4 | 461100 | 1963 | 0.4 |
| *Malus* | | Golden Delicious | Clonal | | | 159872 | 17.8 | | | |
| | | Gala | | | | 196164 | 21.6 | | | |
| | *M. fusca* | | Outcrossing | | | 72859 | 11.4 | | | |
| | *M. sieversii* | | | | | 143655 | 14.9 | | | |
| | *M. sylvestris* | | | | | 149501 | 16.4 | | | |
| *Solanum* | | RH89-39-16 | Clonal | 300318 | 22.1 | | | | | |
| | | Heinz 1706 | Inbred | 7056 | 0.5 | | | | | |
| *Oryza* | | MH63 | Inbred | 6943 | 1.1 | | | | | |

**Table S7.** List of species for phylostratigraphic analysis and the source of their proteomes. Columns show: species name, genome version, and source.

| **Species name** | **Version** | **Source** |
|---|---|---|
| *Porphyridium purpureum* | v1.0 | The Porphyridium purpureum Genome Project (http://cyanophora.rutgers.edu/porphyridium/) |
| *Cyanophora paradoxa* | v1.0 | C. paradoxa v.1.0 (Price et al., 2019) |
| *Chlamydomonas reinhardtii* | v5.6 | Phythozome V12 (https://phytozome.jgi.doe.gov/) |
| *Klebsormidium nitens* | v1.0 | K. nitens NIES-2285 (http://www.plantmorphogenesis.bio.titech.ac.jp/~algae_genome_project/klebsormidium/index.html) |
| *Chara braunii* | v1.0 | ORCAE (http://bioinformatics.psb.ugent.be/orcae/) |
| *Physcomitrium patens* | v3.3 | https://peatmoss.online.uni-marburg.de/downloads/ |
| *Selaginella moellendorffii* | v1.0 | Phythozome V13 (https://phytozome.jgi.doe.gov/) |
| *Salvinia cucullata* | v1.1 | FernBase (www.fernbase.org) |
| *Picea abies* | v1.0 | ConGenIE (Conifer Genome Integrative Explorer) web resource (http://congenie.org) |
| *Amborella trichopoda* | v1.0 | Ensembl plants (plants.ensembl.org) |
| *Oryza sativa* | Nipponbare | RiceSuperPIRdb(http://www.ricesuperpir.com/web/nip) |
| *Solanum lycopersicum* | SL5.0 | Solanaceae Genomics Network (https://solgenomics.net/) |
| each of six *Vitis* samples | | our study |

**Table S8.** RNA-seq resources.

| Accession | Type | Class | Treatment | Label | Run | BioProject | Bases (Mb) | LibLayout |
|---|---|---|---|---|---|---|---|---|
| Pinot Noir | Fruit Development | FrDev1 | B20_1 | FrDev1_B20_1 | SRR1748336 | PRJNA260535 | 1369 | SINGLE |
| | | | B20_2 | FrDev1_B20_2 | SRR1748337 | PRJNA260535 | 1716 | SINGLE |
| | | | B20_3 | FrDev1_B20_3 | SRR1748338 | PRJNA260535 | 1597 | SINGLE |
| | | | B26_1 | FrDev1_B26_1 | SRR1748345 | PRJNA260535 | 1548 | SINGLE |
| | | | B26_2 | FrDev1_B26_2 | SRR1748346 | PRJNA260535 | 1845 | SINGLE |
| | | | B26_3 | FrDev1_B26_3 | SRR1748347 | PRJNA260535 | 1724 | SINGLE |
| | | FrDev2 | VR_1 | FrDev2_VR_1 | ERR3863114 | PRJEB36552 | 2022 | SINGLE |
| | | | VR_2 | FrDev2_VR_2 | ERR3863115 | PRJEB36552 | 1965 | SINGLE |
| | | | VR_3 | FrDev2_VR_3 | ERR3863116 | PRJEB36552 | 1971 | SINGLE |
| | | | MR_1 | FrDev2_MR_1 | ERR3863117 | PRJEB36552 | 1844 | SINGLE |
| | | | MR_2 | FrDev2_MR_2 | ERR3863118 | PRJEB36552 | 1745 | SINGLE |
| | | | MR_3 | FrDev2_MR_3 | ERR3863119 | PRJEB36552 | 2050 | SINGLE |
| | | FrDev3 | inflo_1 | FrDev3_inflo_1 | ERR4606196 | PRJEB39263 | 1658 | SINGLE |
| | | | inflo_2 | FrDev3_inflo_2 | ERR4606197 | PRJEB39263 | 1977 | SINGLE |
| | | | inflo_3 | FrDev3_inflo_3 | ERR4606198 | PRJEB39263 | 1375 | SINGLE |
| | | | berry_1 | FrDev3_berry_1 | ERR4606232 | PRJEB39263 | 1536 | SINGLE |
| | | | berry_2 | FrDev3_berry_2 | ERR4606233 | PRJEB39263 | 1814 | SINGLE |
| | | | berry_3 | FrDev3_berry_3 | ERR4606234 | PRJEB39263 | 1475 | SINGLE |
| | Organ Differentiation | OrganD1 | ripe2_1 | OrganD1_ripe2_1 | SRR5405118 | PRJNA381300 | 1792 | PAIRED |
| | | | ripe1_1 | OrganD1_ripe1_1 | SRR5405119 | PRJNA381300 | 1204 | PAIRED |
| | | | ripe1_2 | OrganD1_ripe1_2 | SRR5405120 | PRJNA381300 | 670 | PAIRED |
| | | | leaf_1 | OrganD1_leaf_1 | SRR5405121 | PRJNA381300 | 1186 | PAIRED |
| | | | leaf_2 | OrganD1_leaf_2 | SRR6320536 | PRJNA381300 | 3316 | PAIRED |
| | | | leaf_3 | OrganD1_leaf_3 | SRR6320537 | PRJNA381300 | 2895 | PAIRED |
| | | | ripe2_2 | OrganD1_ripe2_2 | SRR6320540 | PRJNA381300 | 1535 | PAIRED |
| | | OrganD2 | SB_1 | OrganD2_SB_1 | SRR14696992 | PRJNA373967 | 4267 | PAIRED |
| | | | SB_2 | OrganD2_SB_2 | SRR14696991 | PRJNA373967 | 4161 | PAIRED |
| | | | leaf_1 | OrganD2_leaf_1 | SRR14696961 | PRJNA373967 | 2118 | PAIRED |
| | | | leaf_2 | OrganD2_leaf_2 | SRR14696962 | PRJNA373967 | 2732 | PAIRED |
| | | OrganD3 | leaf_1 | OrganD3_leaf_1 | DRR093294 | PRJDB5807 | 2818 | SINGLE |
| | | | leaf_2 | OrganD3_leaf_2 | DRR093295 | PRJDB5807 | 2533 | SINGLE |
| | | | leaf_3 | OrganD3_leaf_3 | DRR093296 | PRJDB5807 | 2838 | SINGLE |
| | | | stem_1 | OrganD3_stem_1 | DRR093297 | PRJDB5807 | 3040 | SINGLE |
| | | | stem_2 | OrganD3_stem_2 | DRR093298 | PRJDB5807 | 2492 | SINGLE |
| | | | stem_3 | OrganD3_stem_3 | DRR093299 | PRJDB5807 | 2454 | SINGLE |

| | | | | | | | | |
|---|---|---|---|---|---|---|---|---|
| | Stress Response | Stress1 | treat_1 | Stress1_treat_1 | SRR19240845 | PRJNA837346 | 7191 | PAIRED |
| | | | treat_2 | Stress1_treat_2 | SRR19240849 | PRJNA837346 | 6957 | PAIRED |
| | | | treat_3 | Stress1_treat_3 | SRR19240850 | PRJNA837346 | 6972 | PAIRED |
| | | | control1 | Stress1_control_1 | SRR19240851 | PRJNA837346 | 6988 | PAIRED |
| | | | control2 | Stress1_control_2 | SRR19240853 | PRJNA837346 | 6942 | PAIRED |
| | | | control3 | Stress1_control_3 | SRR19240854 | PRJNA837346 | 7031 | PAIRED |
| | | Stress2 | drought_1 | Stress2_drought_1 | SRR6706478 | PRJNA433817 | 7203 | PAIRED |
| | | | drought_2 | Stress2_drought_2 | SRR6706480 | PRJNA433817 | 7627 | PAIRED |
| | | | drought_3 | Stress2_drought_3 | SRR6706481 | PRJNA433817 | 7787 | PAIRED |
| | | | control_1 | Stress2_control_1 | SRR6706482 | PRJNA433817 | 7480 | PAIRED |
| | | | control_2 | Stress2_control_2 | SRR6706484 | PRJNA433817 | 8344 | PAIRED |
| | | | control_3 | Stress2_control_3 | SRR6706485 | PRJNA433817 | 7189 | PAIRED |
| | | Stress3 | WD_1 | Stress3_WD_1 | SRR1709063 | PRJNA268857 | 1766 | SINGLE |
| | | | WD_2 | Stress3_WD_2 | SRR1709064 | PRJNA268857 | 1860 | SINGLE |
| | | | WD_3 | Stress3_WD_3 | SRR1709065 | PRJNA268857 | 1683 | SINGLE |
| | | | WW_1 | Stress3_WW_1 | SRR1709066 | PRJNA268857 | 1241 | SINGLE |
| | | | WW_2 | Stress3_WW_2 | SRR1709067 | PRJNA268857 | 1643 | SINGLE |
| | | | WW_3 | Stress3_WW_3 | SRR1709068 | PRJNA268857 | 1787 | SINGLE |
| | | Stress4 | mock_1 | Stress4_mock_1 | SRR14635675 | PRJNA732451 | 5263 | PAIRED |
| | | | mock_2 | Stress4_mock_2 | SRR14635676 | PRJNA732451 | 5673 | PAIRED |
| | | | mock_3 | Stress4_mock_3 | SRR14635677 | PRJNA732451 | 5416 | PAIRED |
| | | | yeast_1 | Stress4_yeast_1 | SRR14635687 | PRJNA732451 | 5210 | PAIRED |
| | | | yeast_2 | Stress4_yeast_2 | SRR14635688 | PRJNA732451 | 5212 | PAIRED |
| | | | yeast_3 | Stress4_yeast_3 | SRR14635689 | PRJNA732451 | 5348 | PAIRED |
| Chardonnay | Fruit Development | FrDev1 | B20_1 | FrDev1_B20_1 | SRR1748348 | PRJNA260535 | 1437 | SINGLE |
| | | | B20_2 | FrDev1_B20_2 | SRR1748349 | PRJNA260535 | 1801 | SINGLE |
| | | | B20_3 | FrDev1_B20_3 | SRR1748350 | PRJNA260535 | 1612 | SINGLE |
| | | | B26_1 | FrDev1_B26_1 | SRR1748357 | PRJNA260535 | 1901 | SINGLE |
| | | | B26_2 | FrDev1_B26_2 | SRR1748358 | PRJNA260535 | 2113 | SINGLE |
| | | | B26_3 | FrDev1_B26_3 | SRR1748359 | PRJNA260535 | 1390 | SINGLE |
| | Organ Differentiation | OrganD1 | leaf_1 | OrganD1_leaf_1 | SRR13403354 | PRJNA691261 | 7951 | PAIRED |
| | | | leaf_2 | OrganD1_leaf_2 | SRR13403355 | PRJNA691261 | 7308 | PAIRED |
| | | | leaf_3 | OrganD1_leaf_3 | SRR13403356 | PRJNA691261 | 7412 | PAIRED |
| | | | EC_1 | OrganD1_EC_1 | SRR13403357 | PRJNA691261 | 7442 | PAIRED |
| | | | EC_2 | OrganD1_EC_2 | SRR13403358 | PRJNA691261 | 7142 | PAIRED |
| | | | EC_3 | OrganD1_EC_3 | SRR13403359 | PRJNA691261 | 7735 | PAIRED |
| | Stress Response | Stress1 | Freeze_1 | Stress1_Freeze_1 | SRR6026704 | PRJNA402079 | 909 | SINGLE |
| | | | Freeze_2 | Stress1_Freeze_2 | SRR6026740 | PRJNA402079 | 817 | SINGLE |
| | | | Freeze_3 | Stress1_Freeze_3 | SRR6026752 | PRJNA402079 | 650 | SINGLE |

| | | | | | | | | |
|---|---|---|---|---|---|---|---|---|
| | | | Control_1 | Stress1_Control_1 | SRR6026703 | PRJNA402079 | 179 | SINGLE |
| | | | Control_2 | Stress1_Control_2 | SRR6026741 | PRJNA402079 | 297 | SINGLE |
| | | | Control_3 | Stress1_Control_3 | SRR6026753 | PRJNA402079 | 280 | SINGLE |
| | | Stress2 | WD_1 | Stress2_WD_1 | SRR1708660 | PRJNA268857 | 1772 | SINGLE |
| | | | WD_2 | Stress2_WD_2 | SRR1708662 | PRJNA268857 | 1908 | SINGLE |
| | | | WD_3 | Stress2_WD_3 | SRR1708663 | PRJNA268857 | 2644 | SINGLE |
| | | | WW_1 | Stress2_WW_1 | SRR1708899 | PRJNA268857 | 1745 | SINGLE |
| | | | WW_2 | Stress2_WW_2 | SRR1708900 | PRJNA268857 | 1937 | SINGLE |
| | | | WW_3 | Stress2_WW_3 | SRR1708901 | PRJNA268857 | 1843 | SINGLE |
| | | Stress3 | CF_1 | Stress3_CF_1 | ERR3163283 | PRJEB31325 | 467 | SINGLE |
| | | | CF_2 | Stress3_CF_2 | ERR3163284 | PRJEB31325 | 424 | SINGLE |
| | | | CF_3 | Stress3_CF_3 | ERR3163285 | PRJEB31325 | 328 | SINGLE |
| | | | CN_1 | Stress3_CN_1 | ERR3163292 | PRJEB31325 | 555 | SINGLE |
| | | | CN_2 | Stress3_CN_2 | ERR3163293 | PRJEB31325 | 472 | SINGLE |
| | | | CN_3 | Stress3_CN_3 | ERR3163294 | PRJEB31325 | 334 | SINGLE |
| Thompson Seedless | | Stress01 | CT7_1 | Stress01_CT7_1 | SRR1927152 | PRJNA275778 | 4996 | PAIRED |
| | | | CT7_2 | Stress01_CT7_2 | SRR1927156 | PRJNA275778 | 5123 | PAIRED |
| | | | CT7_3 | Stress01_CT7_3 | SRR1927158 | PRJNA275778 | 4715 | PAIRED |
| | | | SHT7_1 | Stress01_SHT7_1 | SRR1998070 | PRJNA275778 | 5826 | PAIRED |
| | | | SHT7_2 | Stress01_SHT7_2 | SRR1998071 | PRJNA275778 | 6605 | PAIRED |
| | | | SHT7_3 | Stress01_SHT7_3 | SRR1998072 | PRJNA275778 | 5898 | PAIRED |
| | | Stress02 | C72-3 | Stress02_C72-3 | SRR24906434 | PRJNA981696 | 6024 | PAIRED |
| | | | C72-2 | Stress02_C72-2 | SRR24906435 | PRJNA981696 | 6085 | PAIRED |
| | | | C72-1 | Stress02_C72-1 | SRR24906436 | PRJNA981696 | 6130 | PAIRED |
| | | | C24-3 | Stress02_C24-3 | SRR24906437 | PRJNA981696 | 6103 | PAIRED |
| | | | C24-1 | Stress02_C24-1 | SRR24906439 | PRJNA981696 | 7097 | PAIRED |
| | | Stress03 | WT_2 | Stress03_WT_2 | SRR5984205 | PRJNA395634 | 7156 | PAIRED |
| | | | WT_1 | Stress03_WT_1 | SRR5984206 | PRJNA395634 | 6830 | PAIRED |
| | | | WT36hpi_3 | Stress03_36hpi_3 | SRR5984204 | PRJNA395634 | 7326 | PAIRED |
| | | | WT36hpi_2 | Stress03_36hpi_2 | SRR5984211 | PRJNA395634 | 6267 | PAIRED |
| | | | WT36hpi_1 | Stress03_36hpi_1 | SRR5984212 | PRJNA395634 | 6735 | PAIRED |
| | Fruit Development | FrDev | D2_1 | FrDev_D2_1 | SRR3880212 | PRJNA328814 | 1895 | PAIRED |
| | | | D2_2 | FrDev_D2_2 | SRR3880214 | PRJNA328814 | 1954 | PAIRED |
| | | | D3_1 | FrDev_D3_1 | SRR3880215 | PRJNA328814 | 2145 | PAIRED |
| | | | D3_2 | FrDev_D3_2 | SRR3880223 | PRJNA328814 | 2115 | PAIRED |
| | | | F4d_1 | Stress1_F4d_1 | SRR24116272 | PRJNA952825 | 6813 | PAIRED |
| | | | F2d_3 | Stress1_F2d_3 | SRR24116273 | PRJNA952825 | 7227 | PAIRED |
| | | | F2d_2 | Stress1_F2d_2 | SRR24116274 | PRJNA952825 | 7080 | PAIRED |
| | Stress Response | Stress1 | F2d_1 | Stress1_F2d_1 | SRR24116275 | PRJNA952825 | 7211 | PAIRED |

| | | | | | | | | |
|---|---|---|---|---|---|---|---|---|
| | | | F1d_3 | Stress1_F1d_3 | SRR24116276 | PRJNA952825 | 6530 | PAIRED |
| | | | F1d_2 | Stress1_F1d_2 | SRR24116277 | PRJNA952825 | 6512 | PAIRED |
| | | | F1d_1 | Stress1_F1d_1 | SRR24116278 | PRJNA952825 | 7455 | PAIRED |
| | | | F0d_3 | Stress1_F0d_3 | SRR24116279 | PRJNA952825 | 6790 | PAIRED |
| | | | F6d_3 | Stress1_F6d_3 | SRR24116280 | PRJNA952825 | 6444 | PAIRED |
| | | | F6d_2 | Stress1_F6d_2 | SRR24116281 | PRJNA952825 | 7111 | PAIRED |
| | | | F6d_1 | Stress1_F6d_1 | SRR24116282 | PRJNA952825 | 7064 | PAIRED |
| | | | F4d_3 | Stress1_F4d_3 | SRR24116283 | PRJNA952825 | 6639 | PAIRED |
| | | | F4d_2 | Stress1_F4d_2 | SRR24116284 | PRJNA952825 | 7052 | PAIRED |
| | | | F0d_2 | Stress1_F0d_2 | SRR24116285 | PRJNA952825 | 6917 | PAIRED |
| | | | F0d_1 | Stress1_F0d_1 | SRR24116286 | PRJNA952825 | 6605 | PAIRED |
| | Fruit Development | FrDev1 | W40_1 | FrDev1_W40_1 | SRR3255928 | PRJNA313243 | 6351 | PAIRED |
| | | | W40_2 | FrDev1_W40_2 | SRR3255929 | PRJNA313243 | 7266 | PAIRED |
| | | | W80_1 | FrDev1_W80_1 | SRR3255930 | PRJNA313243 | 7596 | PAIRED |
| | | | W80_2 | FrDev1_W80_2 | SRR3255931 | PRJNA313243 | 7657 | PAIRED |
| | | | W120_1 | FrDev1_W120_1 | SRR3255932 | PRJNA313243 | 8027 | PAIRED |
| | | | W120_2 | FrDev1_W120_2 | SRR3255933 | PRJNA313243 | 8033 | PAIRED |
| | | | B40_1 | FrDev1_B40_1 | SRR3255934 | PRJNA313243 | 7330 | PAIRED |
| | | | B80_1 | FrDev1_B80_1 | SRR3255935 | PRJNA313243 | 7182 | PAIRED |
| | | | B120_1 | FrDev1_B120_1 | SRR3255936 | PRJNA313243 | 7007 | PAIRED |
| | | | B120_2 | FrDev1_B120_2 | SRR3255937 | PRJNA313243 | 7770 | PAIRED |
| | | | B40_2 | FrDev1_B40_2 | SRR3255926 | PRJNA313243 | 7100 | PAIRED |
| | | | B80_2 | FrDev1_B80_2 | SRR3255927 | PRJNA313243 | 7558 | PAIRED |
| | notreat | notreat | leaf_1 | leaf_1 | This study | This study | 6593 | PAIRED |
| | | | leaf_2 | leaf_2 | This study | This study | 6673 | PAIRED |
| | | | leaf_3 | leaf_3 | This study | This study | 6584 | PAIRED |
| | notreat | notreat | leaf_1 | leaf_1 | This study | This study | 6857 | PAIRED |
| | | | leaf_2 | leaf_2 | This study | This study | 6298 | PAIRED |
| | | | leaf_3 | leaf_3 | This study | This study | 5575 | PAIRED |
| Golden Delicious | Fruit Development | FrDev1 | ripe_2 | FrDev1_ripe_2 | SRR5405142 | PRJNA381300 | 1997 | PAIRED |
| | | | ripe_1 | FrDev1_ripe_1 | SRR5405143 | PRJNA381300 | 1301 | PAIRED |
| | | | leaf_1 | FrDev1_leaf_1 | SRR5405145 | PRJNA381300 | 1416 | PAIRED |
| | | | leaf_2 | FrDev1_leaf_2 | SRR6320561 | PRJNA381300 | 1761 | PAIRED |
| | | | imripe1 | FrDev1_imripe1 | SRR7343308 | PRJNA381300 | 1303 | PAIRED |
| | | | imripe2 | FrDev1_imripe2 | SRR7343309 | PRJNA381300 | 1306 | PAIRED |
| | Stress Response | Stress1 | CK_1 | Stress1_CK_1 | SRR9120847 | PRJNA544573 | 6965 | PAIRED |
| | | | CK_2 | Stress1_CK_2 | SRR9120848 | PRJNA544573 | 7412 | PAIRED |
| | | | N_1 | Stress1_N_1 | SRR9120849 | PRJNA544573 | 6390 | PAIRED |
| | | | N_2 | Stress1_N_2 | SRR9120850 | PRJNA544573 | 6495 | PAIRED |

| | | | | | | | | |
|---|---|---|---|---|---|---|---|---|
| | | Stress2 | WL_1 | Stress2_WL_1 | SRR12533150 | PRJNA658028 | 7173 | PAIRED |
| | | | CK_1 | Stress2_CK_1 | SRR12533151 | PRJNA658028 | 7119 | PAIRED |
| | | | WL_2 | Stress2_WL_2 | SRR12533152 | PRJNA658028 | 9607 | PAIRED |
| | | | CK_2 | Stress2_CK_2 | SRR12533153 | PRJNA658028 | 10035 | PAIRED |
| | | Stress3 | treat_1 | Stress3_treat_1 | SRR12534349 | PRJNA658005 | 7008 | PAIRED |
| | | | Untreated_1 | Stress3_Untreated_1 | SRR12534350 | PRJNA658005 | 8048 | PAIRED |
| | | | treat_2 | Stress3_treat_2 | SRR12534351 | PRJNA658005 | 6627 | PAIRED |
| | | | Untreated_2 | Stress3_Untreated_2 | SRR12534352 | PRJNA658005 | 6565 | PAIRED |
| | | | treat_3 | Stress3_treat_3 | SRR12534353 | PRJNA658005 | 6916 | PAIRED |
| | | | Untreated_3 | Stress3_Untreated_3 | SRR12534354 | PRJNA658005 | 8875 | PAIRED |
| | Organ Differentiation | OrganD1 | Fw_3 | OrganD1_Fw_3 | SRR23362255 | PRJNA929720 | 4408 | PAIRED |
| | | | Fw_2 | OrganD1_Fw_2 | SRR23362256 | PRJNA929720 | 3736 | PAIRED |
| | | | Fw_1 | OrganD1_Fw_1 | SRR23362257 | PRJNA929720 | 4127 | PAIRED |
| | | | pop_1 | OrganD1_pop_1 | SRR23362260 | PRJNA929720 | 4101 | PAIRED |
| | | | pop_3 | OrganD1_pop_3 | SRR23362258 | PRJNA929720 | 4345 | PAIRED |
| | | | pop_2 | OrganD1_pop_2 | SRR23362259 | PRJNA929720 | 4277 | PAIRED |
| | | | bud_3 | OrganD1_bud_3 | SRR23362261 | PRJNA929720 | 3871 | PAIRED |
| | | | bud_2 | OrganD1_bud_2 | SRR23362262 | PRJNA929720 | 4509 | PAIRED |
| | | | bud_1 | OrganD1_bud_1 | SRR23362263 | PRJNA929720 | 3875 | PAIRED |

**Table S9.** Bisulfite-seq resources for four *Vitis* taxa.

| ID | Run | BioProject | tissue | Bases | LibraryLayout |
|---|---|---|---|---|---|
| Pinot Noir | SRR6328774 | PRJNA381300 | leaf | 7029105900 | PAIRED |
| | SRR6328775 | PRJNA381300 | leaf | 39836178000 | PAIRED |
| Chardonnay | SRR13403347 | PRJNA691259 | leaf | 29292988590 | PAIRED |
| | SRR25030780 | PRJNA987409 | leaf2 | 17412069900 | PAIRED |
| *V. piasezkii* | This study | This study | leaf | 20569154920 | PAIRED |
| *V. retordii* | This study | This study | leaf | 15929941028 | PAIRED |
| *V. retordii* | This study | This study | leaf | 14886367685 | PAIRED |

**Dataset S1 (separate file).** Centromere and telomere position for six *Vitis* samples.

**Dataset S2 (separate file).**

S2.1. Percentage of diploid and hemizygous genes with a single exon.

S2.2. Average exon length for diploid and hemizygous genes.

S2.3. Average gene length for diploid and hemizygous genes.

S2.4. Average number of exons for diploid and hemizygous genes.

S2.5. Average phylostrata for hemizygous genes in centromere and telomere.

S2.6. Average phylostrata for diploid genes in centromere and telomere.

**Dataset S2 (separate file).**

S3.1. Proportion of hemizygous and diploid genes overlapping with centromeres.

S3.2. Proportion of hemizygous and diploid genes overlapping with telomeres.

S3.3. Average *Ks* values for diploid and hemizygous genes.

S3.4. Average *Ka/Ks* values for diploid and hemizygous genes.

S3.5. Average gene branch length for diploid and hemizygous genes.

S3.6. Average insertion times for diploid and hemizygous LTRs.

S3.7. Percentage of single-copy genes among diploid and hemizygous genes.

**Dataset S4 (separate file).** Functional analysis using DAVID and the corresponding UniProt names.

**Dataset S5 (separate file).**

S5.1. The proportion and average expression of expressed diploid and hemizygous genes.

S5.2. The proportion and average expression of expressed diploid and hemizygous genes for each tissue.

S5.3. The ratio of average expression between expressed hemizygous and diploid genes.

S5.4. Number of expressed DEGs in total of expressed diploid and hemizygous genes in tissue/treatment comparison.

S5.5. The proportion of DEGs in total of expressed diploid and hemizygous genes overall.

S5.6. A summary of gene numbers that were expressed in specific tissues or treatments.

S5.7. A summary of gene numbers for types categorized as common, stress-specific, organ differentiation-specific, and fruit development-specific.

**Dataset S6 (separate file).** Proportion of tissue/treatment-specific genes.

**Dataset S7 (separate file).**

S7.1. The proportion of expressed diploid and hemizygous genes with/without nearby TEs and their average expression.

S7.2. The proportion of expressed diploid and hemizygous genes with/without nearby TEs and their average expression for each tissue.

S7.3. The proportion of expressed diploid and hemizygous genes with diploid and hemizygous TEs and their average expression.

S7.4. The proportion of expressed diploid and hemizygous genes with diploid and hemizygous TEs and their average expression for each tissue.

**Dataset S8 (separate file).** *P* values for Fig.S11-Fig.S14.

**Dataset S9 (separate file).**

S9.1 The number of mC and unmethylated C.
S9.2. Methylation level for diploid and hemizygous genes.
S9.3. Proportion of methylated genes for diploid and hemizygous genes
